# Scywalker: scalable end-to-end data analysis workflow for nanopore single-cell transcriptome sequencing

**DOI:** 10.1101/2024.02.22.581508

**Authors:** Peter De Rijk, Tijs Watzeels, Fahri Küçükali, Jasper Van Dongen, Júlia Faura, Patrick Willems, Lara De Deyn, Lena Duchateau, Carolin Grones, Thomas Eekhout, Tim De Pooter, Geert Joris, Stephane Rombauts, Bert De Rybel, Rosa Rademakers, Frank Van Breusegem, Mojca Strazisar, Kristel Sleegers, Wouter De Coster

**Affiliations:** Neuromics Support Facility, VIB Center for Molecular Neurology, VIB, Antwerp, Belgium; Department of Biomedical Sciences, University of Antwerp, Antwerp, Belgium; Complex Genetics of Alzheimer’s Disease Group, VIB Center for Molecular Neurology, VIB, Antwerp, Belgium; Applied and Translational Neurogenomics Group, VIB Center for Molecular Neurology, VIB, Antwerp, Belgium; Department of Plant Biotechnology and Bioinformatics, Ghent University, Ghent, Belgium; VIB Center for Plant Systems Biology, VIB, Ghent, Belgium; Department of Biomolecular Medicine, Ghent University, Ghent, Belgium; VIB Center for Medical Biotechnology, VIB, Ghent, Belgium; VIB Single Cell Core, VIB, Ghent/Leuven, Belgium

## Abstract

We introduce *scywalker*, an innovative and scalable package developed to comprehensively analyze long-read nanopore sequencing data of full-length single-cell or single-nuclei cDNA. Existing nanopore single-cell data analysis tools showed severe limitations in handling current data sizes. We developed novel scalable methods for cell barcode demultiplexing and single-cell isoform calling and quantification and incorporated these in an easily deployable package. Scywalker streamlines the entire analysis process, from sequenced fragments in FASTQ format to demultiplexed pseudobulk isoform counts, into a single command suitable for execution on either server or cluster. Scywalker includes data quality control, cell type identification, and an interactive report. Assessment of datasets from the human brain, Arabidopsis leaves, and previously benchmarked data from mixed cell lines, demonstrate excellent correlation with short-read analyses at both the cell-barcoding and gene quantification levels. At the isoform level, we show that scywalker facilitates the direct identification of cell-type-specific expression of novel isoforms.

## Introduction

Single-cell and single-nuclei (collectively called single-cell hereafter) transcriptome sequencing has revolutionized our understanding of processes in health and disease, especially in heterogeneous tissue like the human brain^1^. Alternative isoforms, with variation in start sites, splicing of exons, or transcript ends, are highly prevalent in the transcriptome of complex eukaryotes ^2,3^. However, the most commonly used single-cell short-read sequencing methods only result in data from the 5’ or 3’ ends of the transcripts. Even with short-read sequencing covering the entire gene length, there is no direct observation of full-length transcripts and only a partial reconstruction of the splice diversity.

In contrast, long-read sequencing methods from Oxford Nanopore Technologies (ONT) and PacBio enable the sequencing of full-length transcripts and the identification and quantification of all isoform variations. Bulk long-read transcriptome sequencing consistently leads to the discovery of novel isoforms ^4–6^ but lacks the cell-type resolution and may miss isoforms from lowly abundant cell types. There is, therefore, great value in optimizing long-read single-cell transcriptomic methods. Initial strategies to deal with the higher error rate of long-read sequencing included combining short-read sequencing data from the same library ^7,8^, rolling circle amplification ^9^, or dimer nucleotide blocks for barcodes and UMIs ^10^. Methods not relying on short-read sequencing data arose with decreasing nucleotide-level error rates ^11–13^. These methods were tested on relatively small datasets and, in our experience, did not scale to the large datasets (>10,000 cells per sample) currently produced. We developed scywalker to address this issue, creating a more scalable package for adequate analysis of this rich data source using novel methods and extensive parallelization while improving the accuracy of the results and utility simultaneously. In contrast to existing tools, scywalker can, in one command, provide analysis of multiple samples, generating ready-to-use per-cell expression counts and multi-sample per-cell-type summed pseudobulk counts, allowing easy comparison of gene and isoform expression over multiple samples and cell types.

## Results

### Overview of the scywalker pipeline

Scywalker is an integrated workflow for analyzing nanopore long-read single-cell sequencing data, currently tailored to the 10x Genomics microfluidics platform. Scywalker orchestrates a complete workflow from FASTQ to cell-type demultiplexed gene and isoform discovery and quantification. It consists of three main modules (Fig. 1).

**Fig. 1:**
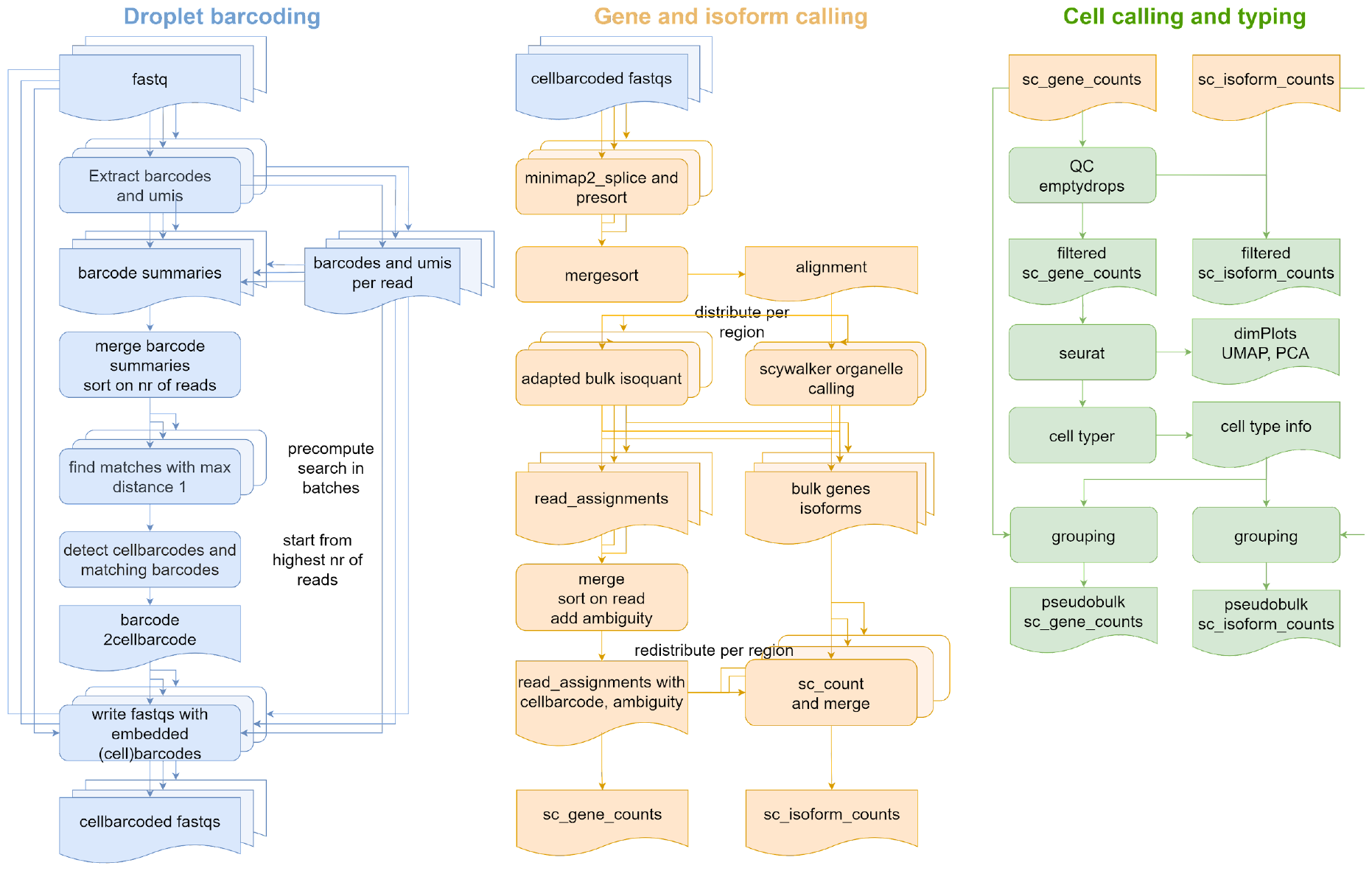
Overview of the scywalker workflow. Parallelized parts are indicated by stacked text boxes. This is the workflow for one sample; multiple samples are always run fully in parallel.

In the first step, 10x droplet barcodes are detected and assigned to all reads using a novel method that does not require a set of given droplet barcodes. All barcodes are sorted according to the number of reads they were found in. The number of reads with a correct barcode can be expected to be substantially higher than those with a barcode containing a specific error. Therefore, the top barcode is considered an accurate droplet barcode, and all barcodes with a single nucleotide difference are searched, corrected, and added to the top barcode. This process is repeated for the remaining barcodes until start barcodes with 20 reads or fewer are reached. The module ends with creating barcoded FASTQ files where each read has its corrected droplet barcode and UMI sequence added.

In the second module, the droplet barcoded reads are aligned to the reference, and isoforms (and genes) are first detected and quantified in a bulk analysis using an adapted version of IsoQuant ^14^ for the non-organelle chromosomes. Gene counting for organelles is done separately using a specific method because organelles, given their different transcription structure and extreme read counts, often pose problems to isoform callers. The IsoQuant and organelle transcript identification produce files specifying the assignment of reads (and thus cell barcodes) to isoforms. This data is used together with the isoform output to produce the quantification of isoforms and genes per droplet barcode in the final step of this module. Based on extra information in the read assignment data (polyA detection, completeness of the match, support for more than one isoform), different types of count are included in the result: weighed (1/n for reads supporting n isoforms), unique (counting only reads uniquely supporting the isoform), strict (only counting unique reads that are 90% complete), and analogous using only reads for which a polyA tail was detected.

Using various approaches initially developed for short-read single-cell analysis, the third module starts by filtering out low-quality and empty droplets from the output of the previous step, and the filtered results are processed using Seurat^15^ for normalization, scaling, and clustering. The cell barcodes are automatically assigned cell types using ScType and scSorter, and finally, pseudobulk counts per cell type are generated based on these assignments. Scywalker can also be used to generate pseudobulk results for user-supplied groupings. When analyzing multiple samples together, scywalker will also make multi-sample, multi-cell type count tables, matching novel isoforms with the same junctions but differences at the ends between different samples.

Scywalker supports scalable parallelization. In order to reduce memory use and improve performance, most steps are subdivided into smaller jobs (indicated by overlapping boxes in Fig. 1), which are efficiently distributed over different processing cores, either on the same computer or over different computers in a cluster. Scywalker is distributed as a portable application directory that contains all dependencies and runs on any Linux system without the need for installation or setting up environments. This facilitates workflow installation and execution, especially on clusters where root access is not always possible, the external network is not necessarily available, and systems may be heterogeneous.

### Scywalker accurately discovers and quantifies cell barcodes

We isolated single nuclei from four adult human brains and performed droplet barcoding and cDNA generation using 10x Chromium, aiming to get ∼10,000 nuclei per sample (hereafter mentioned as brain1-4). The resulting libraries were sequenced both on ONT PromethION (P24) and Illumina NovaSeq 6000 (run details see Supplementary Table 1). The short read sequencing is not needed for the scywalker analysis but provides a reference to evaluate the long read analysis results on the cell and gene count level.

We compared the UMI-corrected read counts per cell barcode found by scywalker in the long reads and Cell Ranger for the short read data. The correlation between the two (Supplementary Fig. 1) is very high (R=0.984, 0.993, 0.995, 0.994 respectively for brain1, brain2, brain3, brain4), showing that scywalker accurately finds droplet barcodes and assigns reads to them.

Furthermore, scywalker also produces accurate results at the gene count per cell level when compared to the short-read data (Fig. 2). While some genes show up only in either short or long-read data here, these are, with a few exceptions, mostly confined to lower expressed ones, and the overall correlation between short and long read data is still very high (R=0.934, 0.932, 0.931, 0.914 respectively for brain1, brain2, brain3, brain4).

**Fig. 2:**
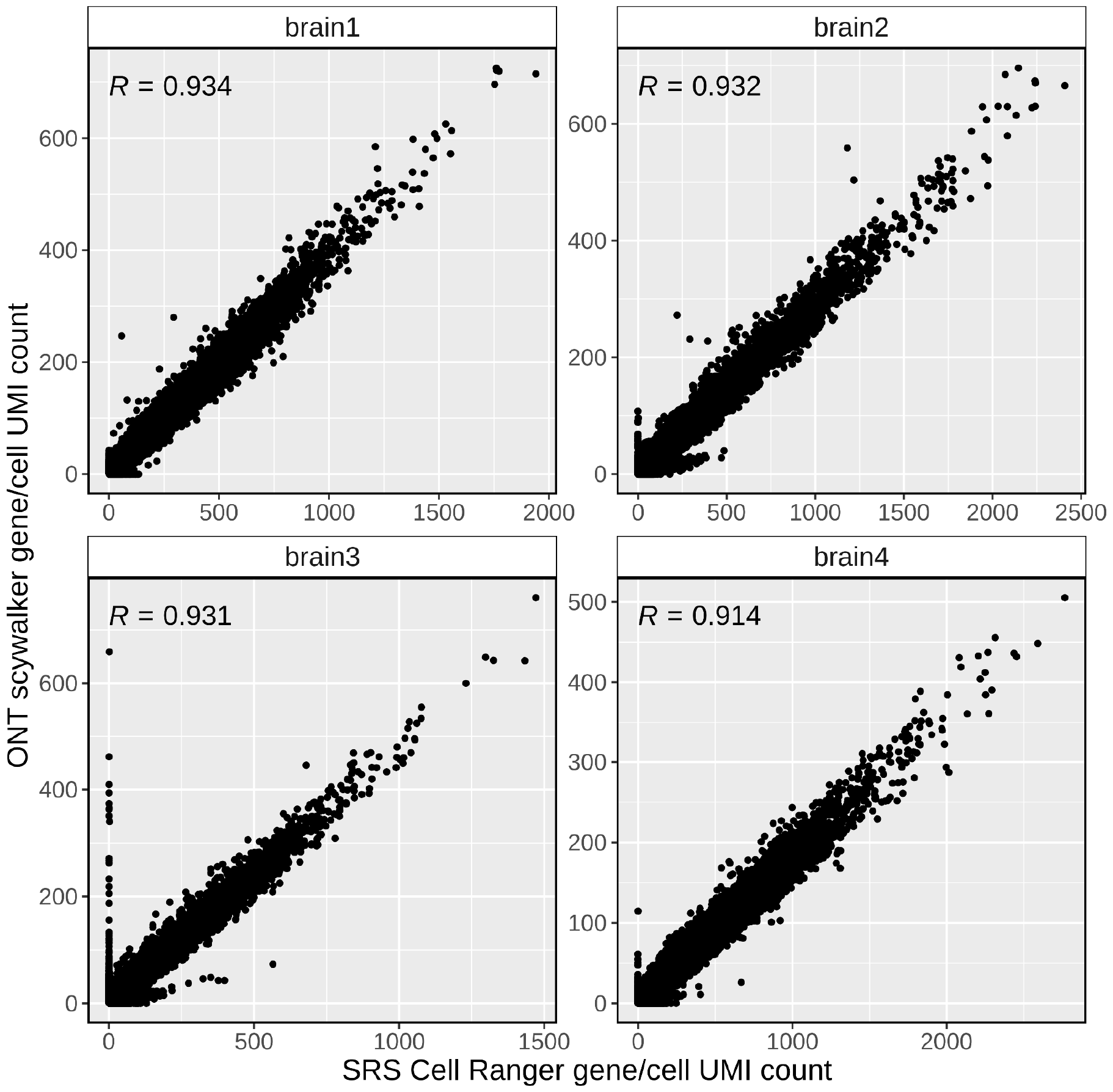
Scywalker UMI counts per gene and cell compared to their respective short-read Cell Ranger results for the four human brain samples. Sample-specific Pearson correlation coefficients (R) are shown on the upper left corners of each panel. *y-axis*, scywalker UMI counts per gene and cell from long-read sequencing data; *x-axis*, Cell Ranger UMI counts per gene and cell from short-read sequencing data. SRS, short-read sequencing; UMI, unique molecular identifier.

### Comparison to other software

While we had successfully tested the wf-single-cell pipeline provided by ONT (formerly known as Sockeye)^13^ on smaller datasets, it could not handle the size of the single-nuclei brain dataset (>10000 nuclei, >100m reads per sample). As an alternative, we tried the combination of BLAZE ^11^ for barcode discovery and FLAMES^12^ for isoform analysis, which did run successfully but took more than a week to complete. To properly compare scywalker to both wf-single-cell and BLAZE-FLAMES and show that scywalker also works on data from less accurate data from earlier iterations of the sequencing chemistry and basecaller versions (Guppy 3.1.5), we downloaded the scmixology2 (GSE154870) data set^16^. We used this smaller dataset (target 183 cells, 22M reads) to benchmark the different packages.

At the cell level (Supplementary Fig. 2), scywalker shows a higher correlation (R=0.942) with the short read data than wf-single-cell (R=0.734) and BLAZE-FLAMES (R=0.65). The largest differences for scywalker were cells detected only in the ONT data, while the two other packages mainly missed cells found in the short read data. Moreover, for the gene counts per cell level (Supplementary Fig. 3), scywalker also shows a higher correlation (R=0.89) than wf-single-cell (R=0.765) and BLAZE-FLAMES (R=0.563).

Scywalker includes downstream analysis of per-cell gene counts, which was successful for this data set. Scywalker found the expected 5 clusters and could assign them to the five cell lines using a custom marker set (Supplementary Fig. 4).

A comparison of the run times for this smaller scmixology2 data set, tested on a system with a 24-core EPYC 7443P and 512G memory, shows comparable results for the three tools, with BLAZE-FLAMES being the fastest (3h19), scywalker second (3h37) and wf-single-cell the slowest (5h41). The analysis of the larger brain1 sample using scywalker took 14h33, or around four times as long, on the same benchmark system. However, the analysis of this larger data set using BLAZE-FLAMES took 172h06, or over 50 times as long as the smaller scmixology2.

The correlation of the cell (Supplementary Fig. 5) and gene quantification (Supplementary Fig. 6) compared with the short-read results was also lower for BLAZE-FLAMES (R=0.888 and R=0.868) than for scywalker (R=0.984 and R=0.934) for the brain1 data set.

### Scywalker enables the detection of differential usage of transcripts between different brain cell types

Scywalker successfully assigned, for the four brain samples, the different brain nuclei cell types (Fig. 3, Supplementary Fig. 7) and generated pseudobulk files for both gene and isoform counts, and combined these in multi-sample, multi-celltype gene and isoform count files.

**Fig. 3:**
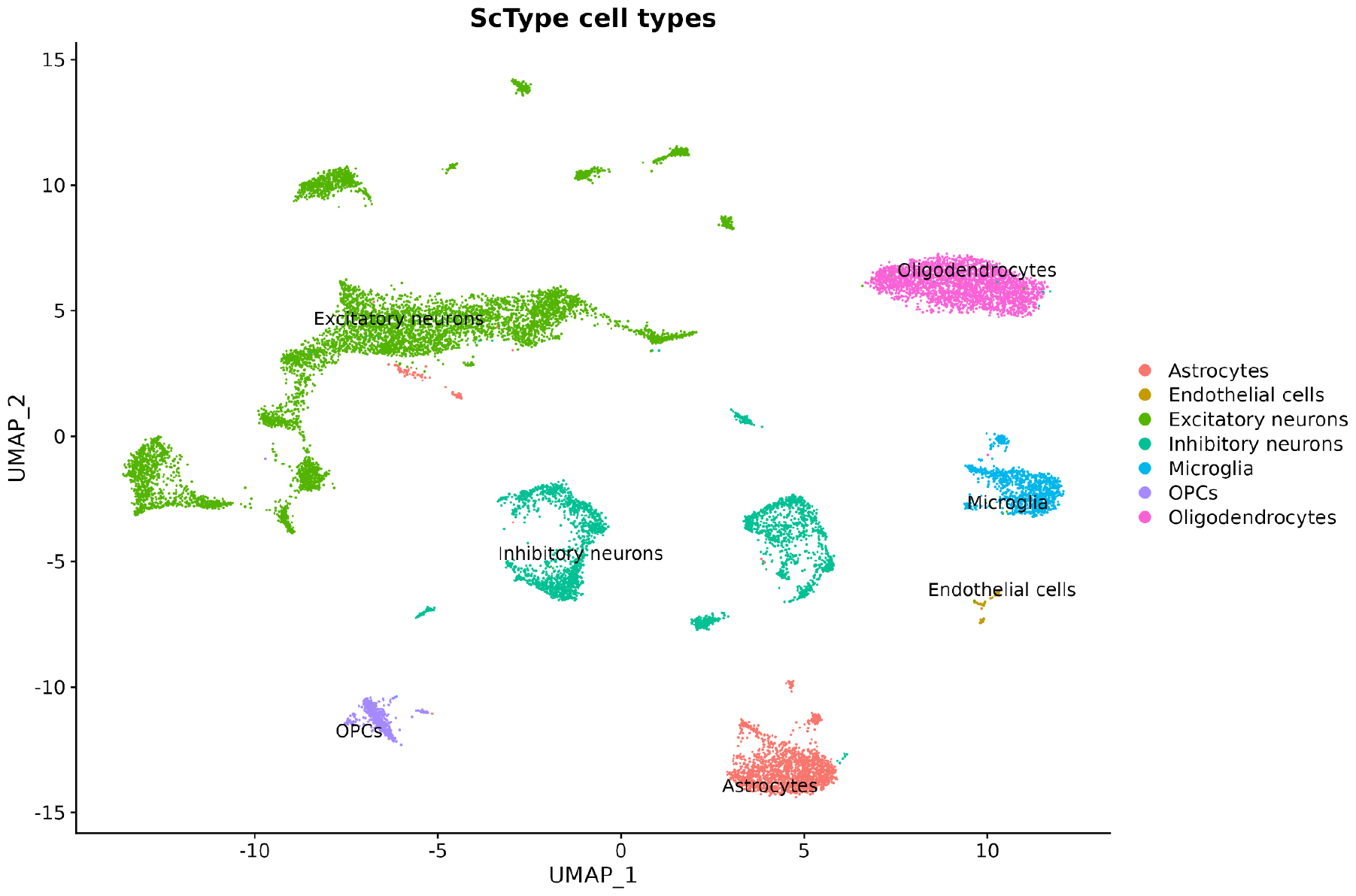
UMAP plot generated by scywalker showing cell-type assignments by ScType in different colors for the brain1 sample.

To evaluate the results of scywalker at the isoform level, we first compared the number of identified isoforms across cell types using the pseudobulk counts from the four brain samples. Excitatory neurons exhibited the highest number of identified isoforms, of which 11.3% were novel (Fig. 4A), followed by inhibitory neurons and oligodendrocytes. Endothelial cells, the cell type with the lowest cell count, also exhibited the fewest identified isoforms (Fig. 4B). Additionally, we observed a correlation between the number of isoforms and the count of novel isoforms (R=0.99, p-value=1.9×10^−5^) (Fig. 4B). Scywalker was also able to identify multiple transcripts per gene, as shown in Fig. 4C. Among the identified isoforms, considered if present in a minimum of 2 samples in at least one cell type (188806 isoforms), ∼40% of them were present in at least five distinct cell types, and 26% were neuron-specific (Fig. 4D).

**Fig. 4:**
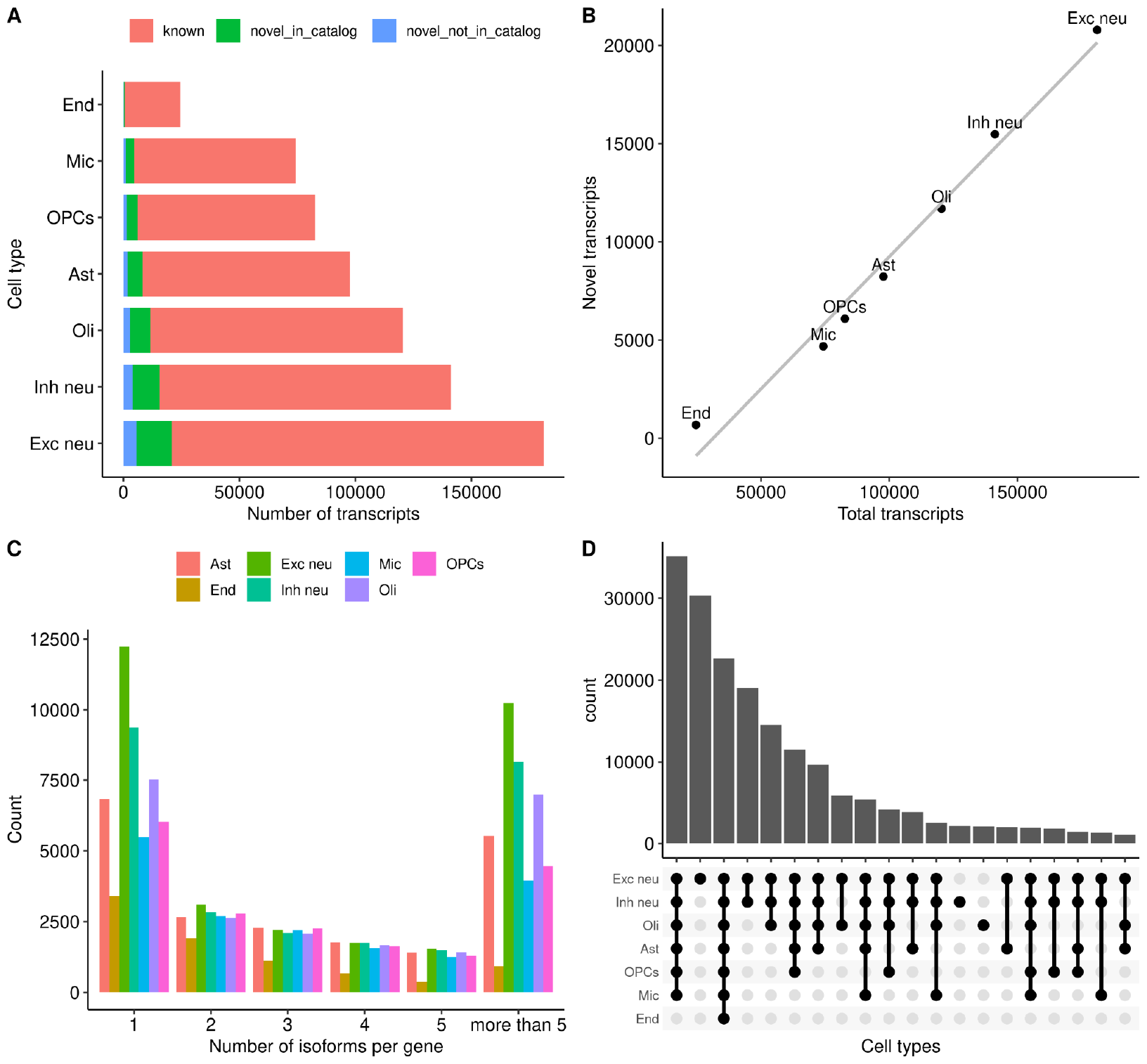
Overview of the scywalker performance at the isoform level. (A) Bar plot representing the number of transcripts identified by cell type. Colors represent different transcript types. (B) Scatter plot correlating the number of total transcripts and novel transcripts (Pearson correlation R=0.99, p=1.9×10^−5^). (C) Bar plot showing the number of isoforms per gene by cell type. (D) Upset plot representing the intersections of identified transcripts per cell type. Exc neu: excitatory neurons; Inh neu: inhibitory neurons; Ast: astrocytes; End: endothelial cells; Oli: oligodendrocytes; OPCs: oligodendrocyte progenitor cells.

Next, we performed a differential transcript usage (DTU) analysis to assess the capability of scywalker to detect isoform expression variation. We compared the proportions of isoforms within each gene between neuronal cell types (excitatory and inhibitory neurons) and glial cell types (oligodendrocytes, astrocytes, OPCs, and microglia). We found 301 genes with DTU (gDTU) (FDR<0.05), allowing the identification of neuron-specific isoforms of specific genes such as *SEPTIN8* (gFDR=6.16×10^−46^, ENST00000378719.7 tFDR=1.09×10^−57^) (Fig. 5A). Septins are known to undergo extensive alternative splicing. In the case of *SEPTIN8*, it was previously suggested that some of its transcripts are ubiquitously expressed across tissues, while others show most expression in the central nervous system^17^. Additionally, in 24 other genes, novel isoforms were found to be differentially used between neurons and glia, for example for *PNKD* (gFDR=2.21×10^−16^, novel transcript tFDR=1.38×10^−9^) (Fig. 5B).

**Fig. 5:**
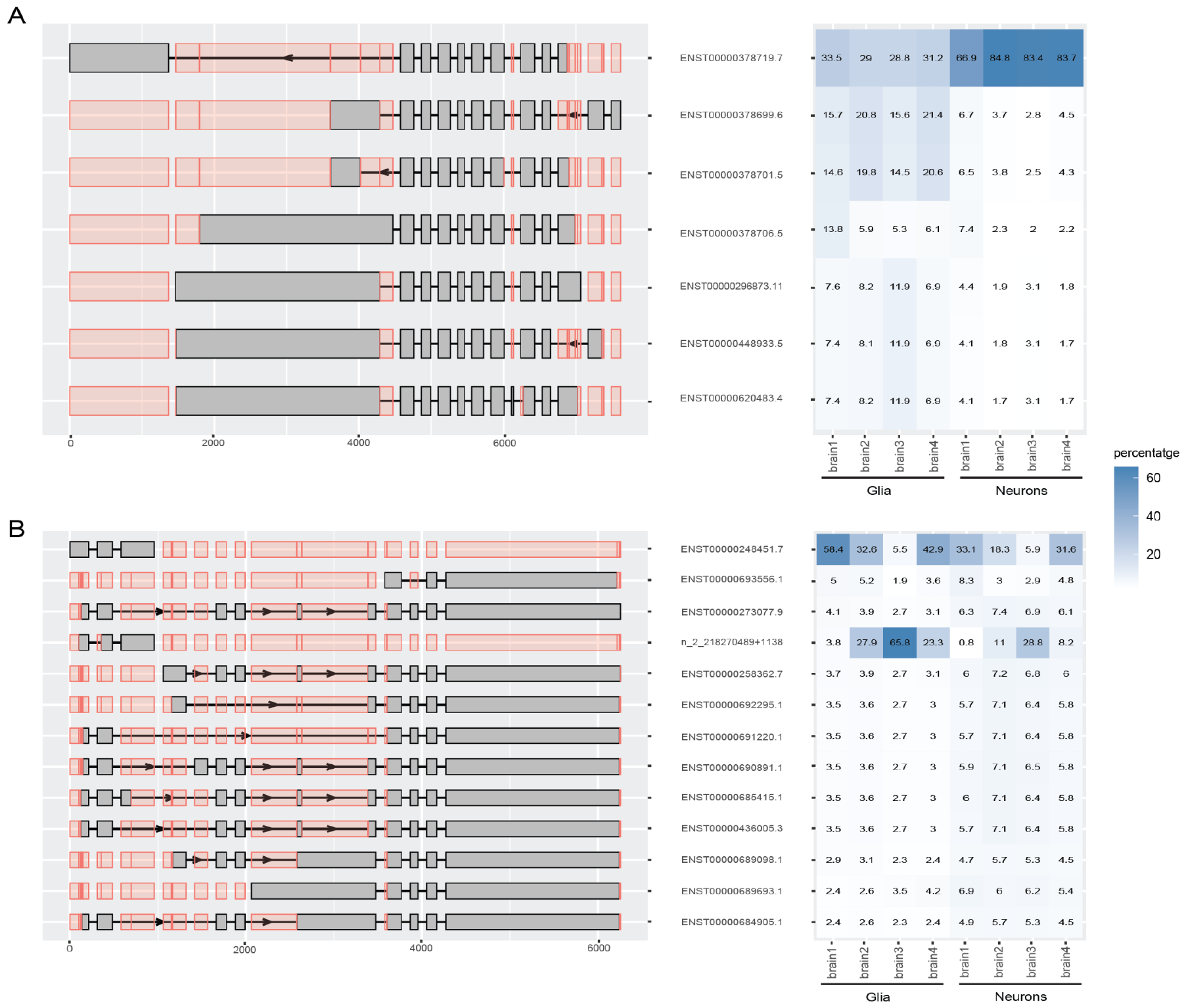
Transcript proportion and exon representation of two gDTUs as generated by scywalker, (A) for *SEPTIN8* and (B) for *PNKD*. In the left panel, all the analyzed isoforms of the gene are shown, with gray rectangles representing the exons of each transcript and red rectangles indicating all potential exon positions for reference. The right panel illustrates the observed percentage abundance of each transcript within groups (glia or neurons) for each brain sample.

### Scywalker analysis is also suitable for non-human data

To showcase its broad applicability in the analysis of non-human species, we also tested scywalker on two large plant single-cell data sets. For this, protoplasts were prepared from mature *Arabidopsis thaliana* leaves, followed by the generation of cell-barcoded cDNA using the 10x Chromium, obtaining ∼10,000 cells per sample. The cDNA was subsequently sequenced on both NovaSeq 6000 and ONT PromethION P24. The short read data (377M reads and 467M reads) was analyzed using Cell Ranger, while the long read data (220M and 230M reads) was analyzed with scywalker. As for the human brain samples, the short read (Cell Ranger) and long read (scywalker) UMI corrected read counts per cell showed excellent correlation (R=0.973 and R=0.988 respectively for plant1 and plant2) (Supplementary Fig. 8).

Similarly, for the plant leaf samples, the UMI gene counts were compared for all genes (except rRNA genes) in cells that had more than a single count in at least one of the two datasets. Here, the correlation between scywalker and Cell Ranger counts is also high (R=0.89 and R=0.898 respectively for plant1 and plant2) (Supplementary Fig. 9), thus demonstrating that scywalker provides reliable single-cell expression metrics in human and plant long-read scRNA-seq samples.

After data preprocessing and cell gene count estimation, scywalker performed follow-up single-cell analysis, including clustering and assignment of cell types. After providing a custom list of cell-type marker genes (Supplementary Table 3), scywalker successfully assigned major leaf cell types (Fig. 6, Supplementary Fig. 10), including the epidermis (Fig. 6B), mesophyll (Fig. 6C), bundle sheath (Fig. 6D) and other leaf cell types.

**Fig. 6:**
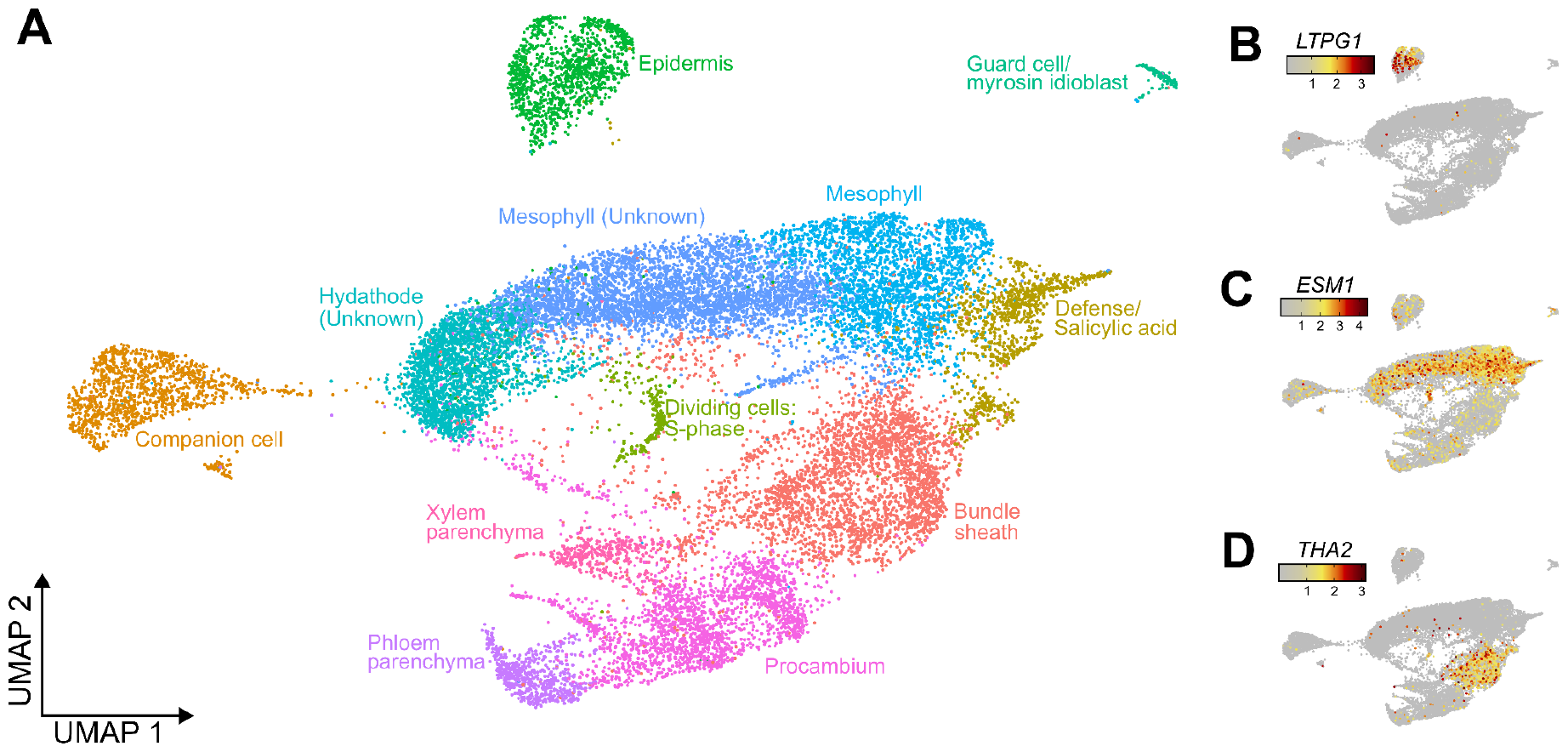
**A**: UMAP plot generated by scywalker showing cell-type assignments by ScType in different colors for plant sample 1. **B-D:** The UMAP plots show the normalized expression of leaf tissue markers for the epidermis (*LTPG1*^18^), mesophyll (*ESM1*^19^), and the bundle sheath (THA2^20^).

At the transcript level, the longer read length produced by ONT allows a more accurate assignment of reads to isoforms and the identification of novel isoforms. Ideally, reads would cover an entire isoform. However, due to various experimental factors (fragmentation, early template switching), this is not the case for most reads (Supplementary Fig. 11). Comparison of plant2 (average read size 967.72) to plant1 (average read size 771.63) clearly illustrates that extracting longer fragments to sequence improves isoform coverage. For plant1, 9.23% of informative reads cover at least 80% of their isoform, rising to 18.63% for plant2.

## Discussion

Long-read single-cell sequencing is essential for a complete transcriptome reconstruction and its cell-type specific alternative isoforms. Compared to short-read sequencing, long reads provide a better resolution at the gene level in organisms with incomplete or inaccurate annotation towards the transcript ends. At the transcript level, more reads are unambiguously assigned to a transcript, offering far better chances of detecting and reconstructing novel isoforms. We sequenced single-nuclei transcriptomes from four human brain samples on the PromethION P24 platform (ONT) as a pilot experiment. In order to obtain a sufficient number of cells from cell types present at lower abundance and to reduce potential sampling errors at that level, we aimed to obtain >10,000 nuclei per sample. The runs generated, on average, >100 million reads per sample. Available analytical workflows did not scale well to this number of cell barcodes and reads. Using novel algorithms and extensive and efficient parallelization, we developed scywalker, which could handle these data readily, analyzing one sample in less than 15 hours on a single 24-core server, and is further scalable to high throughput by running on a cluster. Scalability is important as the field evolves towards datasets assaying more cells at higher depths when the technology matures. The current implementation of the pipeline is tailored to nanopore sequencing of single-cell libraries from the 10X Genomics microfluidics system but is sufficiently flexible to be adapted to accommodate technologies that result in a different read structure if the need arises.

Although long-read sequencing promises to provide more extensive results at the transcript level, short-read Illumina sequencing is the standard for single-cell gene quantification due to its higher per-nucleotide accuracy and yield and its well-established analysis tools. Therefore, we also sequenced the same libraries on an Illumina short-read platform to serve as a reference for benchmarking cell detection and gene quantification in the long-read data. We used the correlation between long-read and short-read data as a measure of the accuracy (even though it is probable that for some genes, the long-read results are more accurate due to less multi-mapping). Using this approach, we showed that, combined with the latest improvements in nanopore sequencing accuracy, scywalker’s algorithms could generate single-cell gene counts that are very highly correlated (R>0.9) with results from short reads and that these can also be successfully used for downstream analyses such as cell typing, thus obviating the need for dual sequencing. Even for the older and less accurate iterations of nanopore sequencing data, the correlation for the scywalker results was high (R=0.89), and the results were usable for downstream analysis. Scywalker also outperformed the other tools (in the instances where they could be tested) on cell and gene level accuracy. In order to ensure the general applicability of scywalker outside of human (or even animal) data, we also generated two plant single-cell data sets (*Arabidopsis thaliana*), which were analyzed successfully by scywalker, also showing very high correlation with their respective short-read results.

Isoform discovery in scywalker is based on IsoQuant, a proven tool showing good results in bulk long-read RNA-seq analysis^14^. In scywalker, the initial discovery is performed without taking cell barcodes into account, i.e., on bulk RNA. This way, isoforms that have low expression in individual cells but are generally present are not discarded due to too low read counts. Based on the read assignments (to isoforms) and their barcodes, gene and isoform counts per cell are generated afterward. Due to experimental issues such as fragmentation, degradation, and premature incorporation of the template-switching oligonucleotide, even in long-read sequencing, most reads do not represent complete isoforms (e.g., Supplementary Fig. 11) and thus can not always be assigned unambiguously to specific transcripts. Scywalker incorporates several ways of dealing with this ambiguity by calculating counts by (i) using reduced weight for ambiguous read assignments, (ii) counting only uniquely assigned reads, (iii) counting only (nearly) complete reads, the latter being e.g., useful for assessing actual support for novel isoforms with different ends.

One of the main goals of long-read sequencing is comparisons at the transcript level, and scywalker provides considerable utility toward that goal. The workflow includes determining cell types, generating pseudobulk data (per cell type), and creating a multi-sample pseudobulk count matrix. As illustrated in the results, this can be used readily for downstream analysis, such as differential isoform usage.

## Conclusion

High-throughput nanopore single-cell sequencing has great potential for uncovering detailed isoform usage across cell types and samples. Scywalker unlocks this potential by providing a highly scalable pipeline capable of handling nanopore samples comprising more than 10,000 cells each and over 100 million reads per sample at volume while simultaneously obtaining excellent accuracy in cell identification and gene quantification. Furthermore, scywalker incorporates automated cell-type assignment, pseudobulk isoform count generation, and sample comparison in this single-command end-to-end workflow. We also demonstrated that the results can be directly used to identify differential transcript usage, including for detecting novel transcripts.

## Methods

### Single-nuclei and single-cell transcriptome sequencing

Four fresh frozen human brain samples (BA10) were provided by the NeuroBiobank of the Born-Bunge Institute (IBB-Neurobiobank), Wilrijk (Antwerp), Belgium; ID: BB190113. The study was approved by the Ethics Committee of the University Hospital Antwerp and the University Antwerp (20/10/107). Nuclei isolation from fresh frozen human brain samples was performed using an adapted density gradient protocol ^21^. A different protease inhibitor (1x cOmplete EDTA-free protease inhibitor) was used, and the lysis time was reduced to 2 minutes.

Arabidopsis (*Arabidopsis thaliana* L. Heynh.) cv. Columbia-0 (Col-0) was grown in a controlled-environment growth chamber (Weiss Technik). For isolation of leaf protoplasts (∼5 excised leaves from four-week-old seedlings pooled per sample), a “tape-sandwich” method was used, where the abaxial epidermis of leaves was peeled using adhesive tape (Scotch® Magic™, 3M) and leaves were immediately immersed in a cell-wall degrading enzymatic buffer in protoplasting buffer (0.6 M mannitol, 20 mM KCl, 10 mM CaCl2, 20 mM MES, 0.1% BSA, 1.0% cellulase R10, 0.3% macerozyme) at pH 5.7. The enzymatic reaction was performed for 1h at room temperature with gentle rotation (30 rpm) in the dark. The solution was filtered through pre-wet 70 μm nylon mesh, collected by centrifugation at 200 × *g* for 6 min, and resuspended in wash buffer (0.6 M mannitol, 20 mM KCl, 10 mM CaCl2, 20 mM MES).

The remainder of the protocol is the same for leaf and brain samples. Beads in emulsion (GEM) generation and droplet barcoding were performed according to the 10x Genomics protocols with Chromium Next GEM Single Cell 3’Kit v3.1 and Chromium Next GEM Chip G Single Cell Kit, aiming for 10,000 cells per sample. Unfragmentated cDNA was prepared for nanopore sequencing on the ONT PromethION (P24) with a combination of the primers from SQK-PCS111, rapid adapters from EXP-RAA114, and auxiliary vials for loading on R10.4.1 flow cells from EXP-AUX003. Data was basecalled on the ONT PromethION (P24) using the Guppy basecaller (v7.0.9) with the SUP model. In parallel, the droplets were further processed for short-read sequencing using the 10x Chromium Next GEM Single Cell 3’ Reagent Kits v3.1 (Dual Index) per the manufacturer’s protocol (10x user guide CG000315). The libraries were sequenced on the Illumina NovaSeq 6000 v1.5 sequencing kit using S4 flow cells, targeting 40,000 reads per cell. The sequencing data was processed with the Cell Ranger pipeline.

### Implementation of the scywalker pipeline

The scywalker pipeline (Fig. 1) is implemented within the GenomeComb framework ^22^, a toolset for the efficient analysis of various types of sequencing data.

#### Scywalker droplet barcoding

The first step of identifying the raw barcodes and UMIs in the reads is distributed per FASTQ file. The adapter fragment in the reads is located using minimap2^23^, with the adapter sequence as the reference and alignment of the reads with settings allowing for short matches (-a --secondary=no -x map-ont -t 4 -n 1 -m 1 -k 5 -w 1 -s 20). The output alignments are then parsed to determine, for each read, which part of the sequenced fragment matches the adapter. By default, the 16 bases after the adapter are taken as the cell barcode (<barcodesize> parameter), and the following 12 bases for the UMI (<umisize> parameter). The results of barcode identification (read name, barcode, UMI) are written to a tab-separated barcode file accompanying the FASTQ.

All barcode files are merged, summarized, and sorted by read count. The top barcode (largest read count) is taken as a correct droplet barcode, and the remaining barcodes are searched for barcodes with one difference using cost-only dynamic programming with a cost cut-off of 1. This search is precomputed in parallel in 50 batches. All hits will be assigned to the initial droplet barcode and removed from the list for further processing. This procedure is then repeated with the following (free) barcode in the list until it has fewer than (by default) 20 reads, or until (by default) 100,000 droplets have already been identified. If a whitelist of all potential droplet barcodes is given, only barcodes in this list will be processed. The resulting mapping of read barcodes to droplet/cell barcodes will be combined with the FASTQs and their barcoding files to generate FASTQs where the corrected (droplet) barcode and UMI are prepended before the read name and added as FASTQ comments.

#### Scywalker Gene and isoform calling

The barcoded FASTQ files are aligned to the reference genome with minimap2 ^23^ using the splice preset in separate jobs for each file and coordinate sorted using a modified version of gnu-sort ^24^. The presorted alignments are merged using the gnu-sort mergesort, resulting in one sorted alignment file in bam format.

Based on this alignment, IsoQuant identifies isoforms as on bulk data, essentially ignoring cell barcodes ^14^. We have made several adaptations to running IsoQuant for the scywalker pipeline. Scywalker processes the read alignments to the genome in smaller regions (default target 5Mbase, but splits can only happen in 250kbase regions without known genes) to improve parallelization and reduce peak memory use. These are run in separate jobs instead of threads (to allow distribution over a cluster). Furthermore, the separate results for known and novel isoforms produced by IsoQuant are integrated. Identifiers of novel isoforms and genes are changed to a unique identifier based on the genomic location to avoid collisions when separately processed regions are merged. Scywalker uses a custom algorithm for analyzing organelle genomes. Only gene locations are loaded into memory to minimize memory use, and numerous read alignments are streamed. For each read alignment, the CIGAR string is parsed to define different alignment regions of the read, and the percent overlap with genes is calculated. If multiple overlaps exist, the non-ribosomal RNA gene with the highest pct overlap is selected. One of the following assignment_events is assigned to the alignment overlapping a gene: mono_exon_match if the alignments start and end coordinates differ a maximum of 3 bases from the known gene location, mono_exon_enclosed for alignments entirely within the gene (but not a match) and mono_exon_overlap for alignments overlapping at least 5% of the gene.

After the regions are processed separately, the read assignment info is concatenated, sorted on the read name (which starts with barcode and UMI info), and processed per block of assignments concerning the same read (has the same barcode and UMI, or the same read name if those were not found). Reads uniquely assigned to one isoform are copied directly into the final read assignment file. Multiple assignments in a block are first filtered: assignments that are less informative than others (significantly shorter, not in a gene, while others match a gene/isoform) in the block are removed. Additional information that will serve as correction factors in the counting step is added to the read assignments: *umicount* (the number of reads having this same UMI), *ambiguity* (the number of isoforms supported by the read), *gambiguity* (the number of genes/locations supported), while information such as *covered_pct* (how much of the isoform is covered by the read) and *inconsistency* (level of inconsistency the alignment shows with the isoform structure) will also be used in providing different types of counts. Finally, the read assignments file is (re)sorted according to genomic location.

Scywalker returns different types of counts as separate fields in the results. These are calculated simultaneously by parsing the read assignments file and keeping a tally for each type of count and cell. A weight is added to the appropriate tallies for each read assignment. For the “weighted” isoform count, a weight of 1/(*ambiguity*\**umicount*) is added, while the “unique” count only takes reads into account that uniquely support one transcript, and “strict” only counts unique reads that cover ≥ 90% of the assigned transcript. Similar results are provided in the “aweighted”, “aunique” and “astrict” columns, but limited to reads with a detected polyA tail. For the default gene “count” a weight of 1/(*gambiguity*\**umicount*) is added, including for reads mapping to introns of the genes (which is also the default for Cell Ranger), while “nicount” only counts reads matching an isoform.

#### Scywalker cell filtering and typing

Droplets that contain nuclei are determined based on the EmptyDrops algorithm ^25^. To improve cell clustering, novel genes detected by scywalker are removed, as well as cells with more than 5% mitochondrial reads. Cell clustering is performed at a relatively low resolution of 0.5, and all principal components are used when computing the nearest neighbor graph and UMAP (Fig. 3). We perform clustering at this resolution to confine the automated assignment of cell types to the identification of the major populations in the sample. At this stage, we only expect to capture high-level populations. By lowering the clustering resolution, we restrict the number of identified clusters to increase the grouping of cells that belong to the same class.

Each cell is assigned to one of the cell types in a user-provided marker file using ScType ^26^ and scSorter ^27^. Users can instead opt to give only the tissue from which the sample originates, in this case, only ScType will be run using its general (human) marker database. The results are tab-separated group files (one for each typer) that link cell to cell type. These are then used together with the per-cell gene and isoform files to generate pseudobulk files giving gene and isoform counts for each cell type (in separate columns). Scywalker will make multi-sample, multi-cell-type count tables when running on a project with multiple samples. Novel isoforms with the same junctions, but differences at the ends from different samples will be merged into one line. The resulting pseudobulk files can be used directly for downstream analysis. Users can also inspect (recommended) and adapt the automatically assigned cell annotations, e.g., a more in-depth annotation with careful expert considerations. After recording adapted cell type annotations in a group file, a user can easily re-run scywalker’s pseudobulk module to obtain pseudobulk matrices for the newly assigned populations.

#### Scywalker reporting

Alignment metrics from the minimap2 bam file are analyzed using cramino in spliced mode^28^, additionally complemented with plots and metrics from identified nuclei and transcript assignment and summarized in an HTML report. The report provides key metrics of the sequencing and cell and gene identification, several interactive plots, such as a knee plot and quality control filters, and information on the read length, including a histogram and a comparison of read lengths against the number of exons detected per read. Furthermore, a histogram shows the number of genes identified per cell and the overlap between reads and known or novel genes, antisense, intergenic, or intronic intervals. Finally, the report includes the UMAP for cell type identification.

### Comparison to existing methods

We compared the scywalker pipeline against the following existing workflows: BLAZE-FLAMES ^11^, respectively versions v1.1.0 and v0.1, and the ONT wf-single-cell ^29^ v0.2.8. Gene level UMI counts for BLAZE-FLAMES were obtained by summing transcript counts per gene. While scywalker outputs a UMI-corrected total cell read count table (independent of the transcript or gene level counts) as well, for accurate comparison of UMI counts per cell between workflows, we considered the cell level UMI counts that were obtained by summing the gene level UMI counts available in all compared workflows. Performance across different workflows was evaluated based on compute time and the correlation of the UMI counts per cell and per gene compared against the short-read quantification. For these correlation comparisons, we only considered the UMI counts per cell and per gene if they were higher than 1 in either of the compared workflows. Moreover, we only included the genes commonly available in the gene annotation references (i.e., GTF/GFF3 files) of the compared workflows, excluding rRNA genes from the comparisons for the plant samples. All software was run using standard parameters (code in Supplementary code), except a maximum edit distance (MAX_DIST) of 1 was used instead of the advised MAX_DIST=2 setting in match_cell_barcode step of the FLAMES workflow for the brain sample because of the higher accuracy of sequencing. We also tested with MAX_DIST=2, which resulted in a substantially lower correlation both for the cell UMI counts comparison (R=0.287 vs R=0.888, Supplementary Fig. 5) and for the gene/cell UMI counts comparison (R=0.644 vs R=0.868, Supplementary Fig. 6). The correlation plots were generated in R (v4.2.0) using the following packages: *ggplot2*^30^, *ggpairs* (as implemented in *GGally*^31^), and *ggpubr*^32^.

### Differential transcript usage between different brain cell types

#### Annotation of brain cell types

Scywalker’s single-cell module creates a Seurat object using the retained cells after EmptyDrops filtering. It then follows the standard Seurat processing workflow with default parameters, with the exception of a slightly lower resolution of 0.5 during clustering. Clusters were then annotated using scSorter and ScType, with the human brain marker genes for the major cell populations (excitatory neurons, inhibitory neurons, astrocytes, microglia, oligodendrocytes, OPCs, endothelial cells, and pericytes, Supplementary Table 2). These markers were selected from a list of literature-based genes based on their segregating capacity, which was determined using Garnett^33^. Markers were only used for annotation if they had an ambiguity of less than 3% in the human brain dataset. Counts were summed per cell population based on these annotations.

#### Differential transcript usage analysis between neuronal and glial cells

We used the resulting pseudobulk counts (*weighted*) as a metric to assess scywalker’s performance at the transcript level. Only transcripts detected in at least half of the samples within a given cell type were retained for subsequent analysis. Transcript counts of the four studied samples were summed for each cell type. The plots to visualize the performance were generated using R (v4.3.2) and the following packages: *ggplot2*^30^, *ggpubr*^32^, and *ggupset* ^30^.

DTU analysis was conducted using the *DRIMSeq* R package ^34^, implementing a Dirichlet-multinomial model. We compared neuronal and glial cell types, with the analysis adjusted for individual variations and conditions (Alzheimer’s Disease or neurologically normal). A two-stage statistical procedure with *stageR* ^35^ accounted for multiple testing. In brief, the DRIMSeq gene p-values were analyzed in a screening stage to determine which genes show signs of DTU (gDTU). In gDTUs, the *DRIMSeq* transcript-level p-values were individually tested for DTU in a confirmation stage, resulting in a corrected p-value for each gene and each transcript of a significant gene. The differential transcript usage plots for specific genes were made using the viz_transcripts tool included with scywalker, which is based on *ggtranscript*^36^ and *ggplot2*^30^.

## Data availability

The human brain data set will be made available in the European Genome-Phenome Archive (EGA), the VIB Data Access Committee will control access to this data. The Arabidopsis sequencing data is available at EBI ArrayExpress under the accession number E-MTAB-13866. The scmixology2 (GSE154870) data set^16^ was obtained from the Sequence Read Archive, under SRR12282457 for the short read data and SRR12282458 for the long read data.

## Code Availability

Scywalker is available on https://github.com/derijkp/scywalker under the GNU General Public License (GPL). The code for the analyses with the different software tools is in the supplementary data.

## Acknowledgments

This work was partly funded by the VIB (Flanders Institute for Biotechnology, Belgium), the University of Antwerp, the Alzheimer’s Association (AARG-20-683760), the Alzheimer Research Foundation Flanders (SAO-FRA 2021/0032), the Fund for Scientific Research Flanders (FWO G030718N), the Centres of Excellence in Neurodegeneration (CoEN6005), and the legacy Maesschalk. PW and WDC are recipients of a postdoctoral fellowship from FWO [12T1722N and 12ASR24N, respectively). LDD receives a PhD fellowship (BOF DOCPRO 44697), and FK a postdoctoral fellowship (BOF 49758) from the University of Antwerp Research Fund. JF receives a Holloway Postdoctoral Fellowship [2022-001] from the Association for Frontotemporal Degeneration (AFTD). This work was supported by the European Research Council (ERC StG TORPEDO; 714055 and ERC CoG PIPELINES; 101043257) to BDR and TE, and by an FWO project grant to FVB (FWO G007723N). Oxford Nanopore Technologies supported this work by providing free PromethION flow cells for sequencing the single-cell Arabidopsis leaf transcriptomes. The authors acknowledge Robin Pottie and Jolien De Block for excellent technical assistance and Kai Xun Chan for discussions. The authors acknowledge Hayden Christensen, Carrie Fisher, and Mark Hamill for inspiring the project. The authors acknowledge the NeuroBiobank of the Born-Bunge Institute (IBB-Neurobiobank), Wilrijk (Antwerp), Belgium; ID: BB190113 for providing the brain samples used in this work. We are thankful to the VIB Single Cell Core and VIB Nucleomics for support and access to the instrument park (vib.be/technologies/).

## Author contributions

PDR, MS, WDC, RR, and KS conceived the project. PDR devised the algorithms and designed and implemented most of the software. TW implemented cell filtering and typing, and WDC implemented the report. PDR, FK, and JVD performed benchmarking. PDR, TW, and JVD did testing. FK performed comparisons of the different workflows. PW analyzed the plant results. TW and JF analyzed differential transcript usage. JVD, LDD, and LD generated the brain single-nuclei libraries for short and long-read sequencing, while CG, TE, and BDR generated the plant data, with SR and FVB coordinating plant data generation and analysis. TDP and GJ performed all nanopore sequencing, coordinated by MS. MS also coordinated software development. KS coordinated the brain data generation and analysis, acquired funding, and administered the project. WDC coordinated data analysis and writing of the manuscript and administered the project. PDR, WDC, TW, FK, JF, and PW drafted the manuscript with input from MS and LDD. All authors reviewed and provided feedback on the manuscript.

