## Supplementary data for "Scywalker: scalable end-to-end data analysis workflow for nanopore single-cell transcriptome sequencing"

#### Code

Code code_comparison: code used to run scywalker, BLAZE-FLAMES, and wf-single-cell on the scmixology2 data set

```
# scmixology2_ont scywalker benchmark run
# =======================================

cd /tmp/benchmark-scmixology2
/usr/bin/time -o time.txt -v \
    scywalker -stack 1 -v 2 -d 24 \
    -sc_expectedcells 183 \
    -cellmarkerfile ../../markers.tsv \
    -threads 4 \
    -refdir /complgen/refseq/hg38 \
    /tmp/benchmark-scmixology2/samples/scmixology2_ont \
    >& startup_process_sample.log

# scmixology2 wf-single-cell benchmark run
# ========================================
cd ~/benchmarks/
mkdir -p /tmp/benchmark-scmixology2/samples/scmixology2_wf/FASTQ
cd /tmp/benchmark-scmixology2/samples/scmixology2_wf
zcat ../../FASTQ/*.FASTQ.gz | gzip > FASTQ/scmixology2_wf.FASTQ.gz

export NXF_SINGULARITY_CACHEDIR=/tmp/singularity

echo 'executor {
      $local {
      cpus = 24
      memory = "500 GB"
      }
}' > my-config.cfg

/usr/bin/time -o time.txt -v \
    nextflow run epi2me-labs/wf-single-cell \
    -r v0.2.8 \
    -profile singularity \
    -c my-config.cfg \
    -w wf-single-cell_workspace \
    --ref_genome_dir /complgen/bin/cellranger-7.1.0/refs/refdata-gex-GRCh38-2020-A \
    --FASTQ FASTQ \
    --sample scmixology2_wf \
    --kit_name 3prime \
```

```
    --kit_version v3 \
    --expected_cells 183 \
    --out_dir wf-single-cell \
    >& wf-single-cell.log

# scmixology2 BLAZE-FLAMES benchmark run
# ======================================
# install
# -------
cd /tmp/benchmark-scmixology2/samples/scmixology2_fm
export mambaversion=22.11.1-4
curl -L -O
"https://github.com/conda-forge/miniforge/releases/download/$mambaversion/Mambaforge
-$mambaversion-Linux-x86_64.sh"
unset PYTHONPATH
rm -rf /home/data/mambaforge
bash Mambaforge-$mambaversion-Linux-x86_64.sh -b

PATH=/home/data/mambaforge/bin:$PATH
mamba init bash
. ~/.bash_profile

wget https://github.com/shimlab/BLAZE/releases/download/v1.1.0/BLAZE_v1.1.0.zip
unzip BLAZE_v1.1.0.zip
cd BLAZE
mamba env remove -n blaze
conda config --set channel_priority false
mamba env create -f conda_env/environment.yml

mamba env remove -n flames
mamba create -y -n flames \
      python=3.7 samtools pysam minimap2 numpy editdistance \
      -c bioconda -c conda-forge
mamba activate flames
git clone https://github.com/LuyiTian/FLAMES.git
cd /tmp/benchmark-scmixology2/samples/scmixology2_fm/FLAMES/src
g++ -std=c++11 -lz -O2 -o match_cell_barcode ssw/ssw_cpp.cpp ssw/ssw.c
match_cell_barcode.cpp kseq.h edit_dist.cpp


# run
# ---
cd /tmp/benchmark-scmixology2/samples/scmixology2_fm
PATH=/home/data/mambaforge/bin:$PATH
mamba init bash
. ~/.bash_profile

#### BLAZE
mamba activate blaze
export
PATH=/home/data/mambaforge/envs/blaze/bin:/home/data/mambaforge/condabin:$PATH
```

```
/usr/bin/time -o time_blaze.txt -v \
    python BLAZE/bin/blaze.py --expect-cells=183 --kit-version=v3 --threads=24 \
    FASTQ
mamba deactivate

#### FLAMES
mamba activate flames
/usr/bin/time -o time_match_cell_barcode.txt -v \
    FLAMES/src/match_cell_barcode FASTQ barcode_statistics.tsv barcoded.FASTQ.gz
whitelist.csv 2 12

/usr/bin/time -o time_flames.txt -v \
       python FLAMES/python/sc_long_pipeline.py \
       --infq barcoded.FASTQ.gz \
       --outdir FLAMES_output \
       --genomefa
/complgen/bin/cellranger-7.1.0/refs/refdata-gex-GRCh38-2020-A/fasta/genome.fa \
       --gff3
/complgen/bin/cellranger-7.1.0/refs/refdata-gex-GRCh38-2020-A/genes/genes.gtf \
       --config_file FLAMES/python/config_sclr_nanopore_default.json

mamba deactivate

# brain1_ont scywalker benchmark run
# ==================================
cd /tmp/benchmark-brain1/samples/brain1_ont
/usr/bin/time -o time.txt -v \
       scywalker -stack 1 -v 2 -d 24 \
       -sc_expectedcells 15000 \
       -cellmarkerfile Cell_markers_Tijs_Jan2024.tsv \
       -threads 4 \
       -refdir /complgen/refseq/hg38 \
       /tmp/benchmark-brain1/samples/brain1_ont \
       >& startup_process_sample.log

# brain1_fm BLAZE-FLAMES benchmark run
# ====================================
cd /tmp/benchmark-brain1/samples/brain1_fm

#### BLAZE
mamba activate blaze
/usr/bin/time -o time_blaze.txt -v \
    python BLAZE/bin/blaze.py --expect-cells=15000 --kit-version=v3 --threads=24 \
    fastq
mamba deactivate

#### FLAMES
mamba activate flames
/usr/bin/time -o time_match_cell_barcode.txt -v \
       FLAMES/src/match_cell_barcode fastq barcode_statistics.tsv barcoded.fastq.gz
whitelist.csv 1 12

/usr/bin/time -o time_flames.txt -v \
       python FLAMES/python/sc_long_pipeline.py \
              --infq barcoded.fastq.gz \
```

```
             --outdir FLAMES_output \
             --genomefa
/complgen/bin/cellranger-7.1.0/refs/refdata-gex-GRCh38-2020-A/fasta/genome.fa \
             --gff3
/complgen/bin/cellranger-7.1.0/refs/refdata-gex-GRCh38-2020-A/genes/genes.gtf \
             --config_file FLAMES/python/config_sclr_nanopore_default.json

mamba deactivate

# Make comparison files
# ---------------------
src=FLAMES_output
sample=brain1_fm
sw csv2tsv $src/transcript_count.csv.gz \
    | sw keyvalue -idfields 'transcript_id gene_id' -keyname cell -valuename count \
    | sw select -f 'transcript=$transcript_id geneid=$gene_id cell count' \
    | sw zst \
    > sc_isoform_counts_filtered-$sample.tsv.zst
sw select -optimization memory -g 'geneid * cell *' -gc 'sum(count)'
sc_isoform_counts_filtered-$sample.tsv.zst \
    | sw select -f 'geneid cell count=$sum_count' \
    | sw zst \
    > sc_gene_counts_filtered-$sample.tsv.zst
sw select -g 'cell *' -gc 'sum(count)' sc_gene_counts_filtered-$sample.tsv.zst \
    | sw select -f 'cellbarcode=$cell count=$sum_count' \
    | sw zst \
    > umis_per_cell_filtered-$sample.tsv.zst
```

### Supplementary Tables

Supplementary Table 1: overview data sets used in this study

| sample | Number of reads | average read size | number of megabases | source |
|---|---|---|---|---|
| brain1 | 109,533,124.00 | 837.36 | 91,719.00 | |
| brain2 | 92,847,419.00 | 795.76 | 73,883.92 | |
| brain3 | 114,379,763.00 | 592.52 | 67,772.50 | |
| brain4 | 93,664,035.00 | 786.75 | 73,689.71 | |
| brain1_srs | 415,477,694.00 | 90.00 | 37,392.99 | |
| brain2_srs | 364,326,042.00 | 90.00 | 32,789.34 | |
| brain3_srs | 266,656,691.00 | 90.00 | 23,999.10 | |
| brain4_srs | 653,867,678.00 | 90.00 | 58,848.09 | |
| scmixology2 | 25,517,285.00 | 1,120.41 | 28,589.86 | SRR12282458 |
| scmixology2_srs | 107,473,860.00 | 91.00 | 9,780.12 | SRR12282457 |
| plant1 | 220,703,149.00 | 771.63 | 170,301.49 | |
| plant2 | 230,532,473.00 | 967.72 | 223,091.72 | |
| plant1_srs | 377,557,437.00 | 90.00 | 33,980.17 | |
| plant2_srs | 467,199,393.00 | 90.00 | 42,047.95 | |

Supplementary Table 2 marker genes for human brain

| geneid | marker | celltype |
|---|---|---|
| ENSG00000067715.15 | SYT1 | Excitatory neurons |
| ENSG00000154146.13 | NRGN | Excitatory neurons |
| ENSG00000091664.9 | SLC17A6 | Excitatory neurons |
| ENSG00000104888.10 | SLC17A7 | Excitatory neurons |
| ENSG00000070808.17 | CAMK2A | Excitatory neurons |
| ENSG00000067715.15 | SYT1 | Inhibitory neurons |
| ENSG00000128683.14 | GAD1 | Inhibitory neurons |
| ENSG00000136750.13 | GAD2 | Inhibitory neurons |
| ENSG00000171885.18 | AQP4 | Astrocytes |
| ENSG00000110436.13 | SLC1A2 | Astrocytes |
| ENSG00000131095.14 | GFAP | Astrocytes |
| ENSG00000129244.9 | ATP1B2 | Astrocytes |
| ENSG00000080493.19 | SLC4A4 | Astrocytes |
| ENSG00000165795.25 | NDRG2 | Astrocytes |
| ENSG00000125398.8 | SOX9 | Astrocytes |
| ENSG00000182578.14 | CSF1R | Microglia |
| ENSG00000169896.18 | ITGAM | Microglia |
| ENSG00000169313.10 | P2RY12 | Microglia |
| ENSG00000019582.17 | CD74 | Microglia |
| ENSG00000168329.14 | CX3CR1 | Microglia |
| ENSG00000125730.18 | C3 | Microglia |
| ENSG00000107099.18 | DOCK8 | Microglia |
| ENSG00000168314.19 | MOBP | Oligodendrocytes |
| ENSG00000197971.16 | MBP | Oligodendrocytes |
| ENSG00000123560.14 | PLP1 | Oligodendrocytes |
| ENSG00000134853.12 | PDGFRA | OPCs |
| ENSG00000173546.7 | CSPG4 | OPCs |
| ENSG00000038427.16 | VCAN | OPCs |
| ENSG00000184113.10 | CLDN5 | Endothelial cells |
| ENSG00000261371.6 | PECAM1 | Endothelial cells |
| ENSG00000169908.12 | TM4SF1 | Endothelial cells |
| ENSG00000102755.12 | FLT1 | Endothelial cells |
| ENSG00000179776.19 | CDH5 | Endothelial cells |
| ENSG00000166825.15 | ANPEP | Pericytes |

Supplementary Table 3 marker genes for Arabidopsis leaf

| geneid | marker | celltype |
|---|---|---|
| AT1G77990 | SULTR2;2 | Bundle sheath |
| AT3G04520 | THA2 | Bundle sheath |
| AT5G57350 | AHA3 | Companion cell |
| AT1G79430 | APL | Companion cell |
| AT1G22710 | SUC2 | Companion cell |
| AT3G22231 | PCC1 | Defense/Salicylic acid |
| AT2G14560 | LURP1 | Defense/Salicylic acid |
| AT5G03350 | AT5G03350 | Defense/Salicylic acid |
| AT1G76540 | CDKB2-1 | Dividing cells: G2/M-phase |
| AT1G44110 | CYCA1-1 | Dividing cells: G2/M-phase |
| AT2G28740 | HIS4 | Dividing cells: S-phase |
| AT3G27060 | TSO2 | Dividing cells: S-phase |
| AT1G07370 | PCNA1 | Dividing cells: S-phase |
| AT1G27950 | LTPG1 | Epidermis |
| AT4G21750 | ATML1 | Epidermis |
| AT1G01120 | KCS1 | Epidermis |
| AT1G54040 | ESP | Epidermis |
| AT3G24140 | FMA | Guard cell/myosin idioblast |
| AT5G26000 | TGG1 | Guard cell/myosin idioblast |
| AT5G25980 | TGG2 | Guard cell/myosin idioblast |
| AT1G08810 | MYB60 | Guard cell/myosin idioblast |
| AT3G16660 | AT3G16660 | Hydathode |
| AT3G16670 | AT3G16670 | Hydathode |
| AT3G14210 | ESM1 | Mesophyll |
| AT1G70760 | NdhL | Mesophyll |
| AT4G12970 | EPFL9 | Mesophyll |
| AT3G27690 | LHCB2.4 | Mesophyll |
| AT3G48740 | SWEET11 | Phloem parenchyma |
| AT5G23660 | SWEET12 | Phloem parenchyma |
| AT5G61480 | PXY | Procambium |
| AT3G15990 | SULT3 4 | Procambium |
| AT2G36120 | DOT1 | Procambium |
| AT3G01680 | SEOR1 | Sieve element |
| AT3G01670 | SEOR2 | Sieve element |
| AT5G19530 | ACL5 | Xylem parenchyma |
| AT4G32880 | ATHB-8 | Xylem parenchyma |
| AT5G60490 | FLA12 | Xylem parenchyma |

### Supplementary Figures

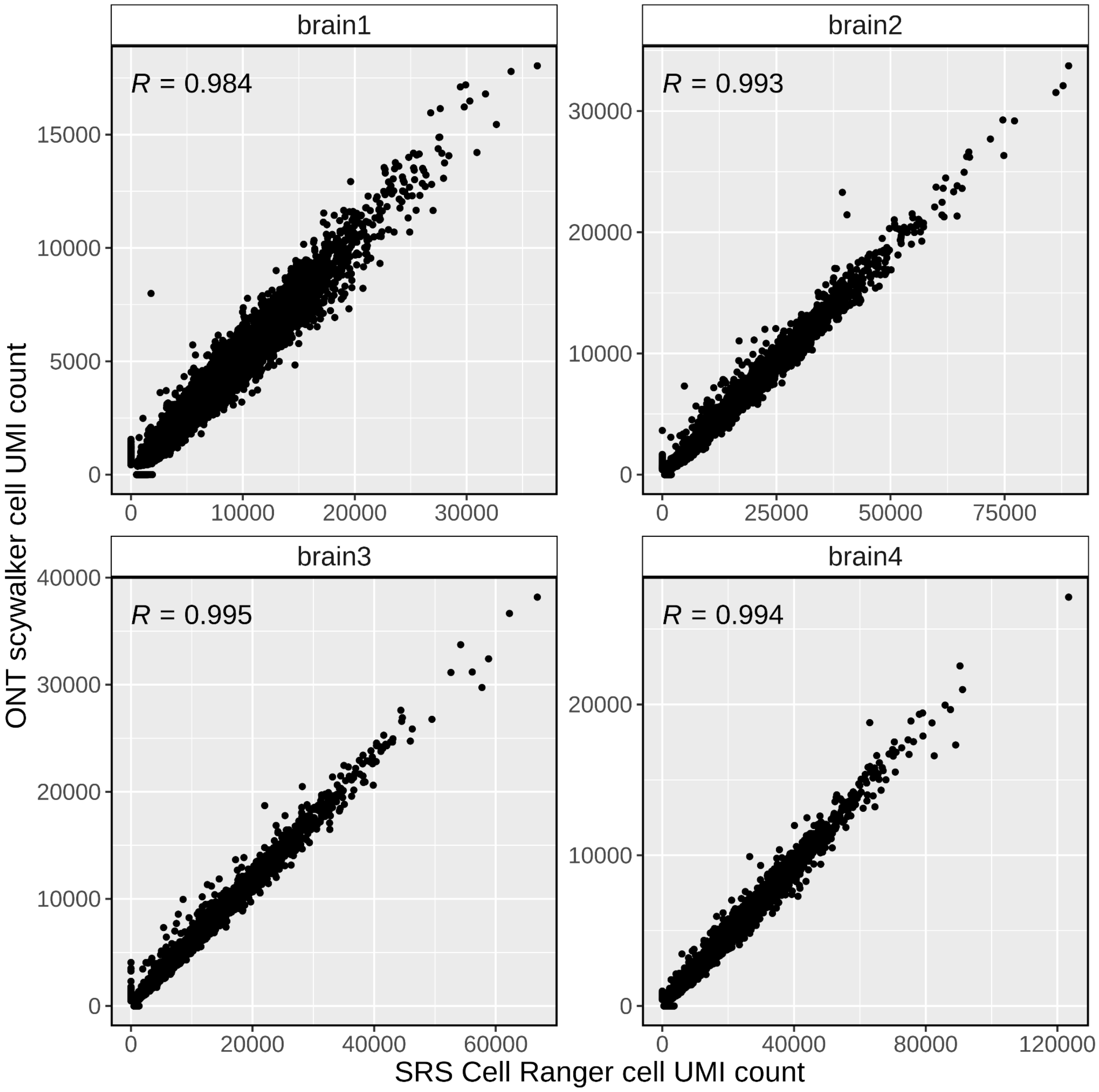


Supplementary Fig. 1: Scywalker UMI counts per cell compared to their respective short read Cell Ranger results for the four human brain samples. Sample-specific Pearson correlation coefficients (R) are shown on the upper left corners of each panel. *y-axis*, scywalker UMI counts per cell based on long-read sequencing data; *x-axis*, Cell Ranger UMI counts per cell based on short-read sequencing data. SRS, short-read sequencing; UMI, unique molecular identifier.

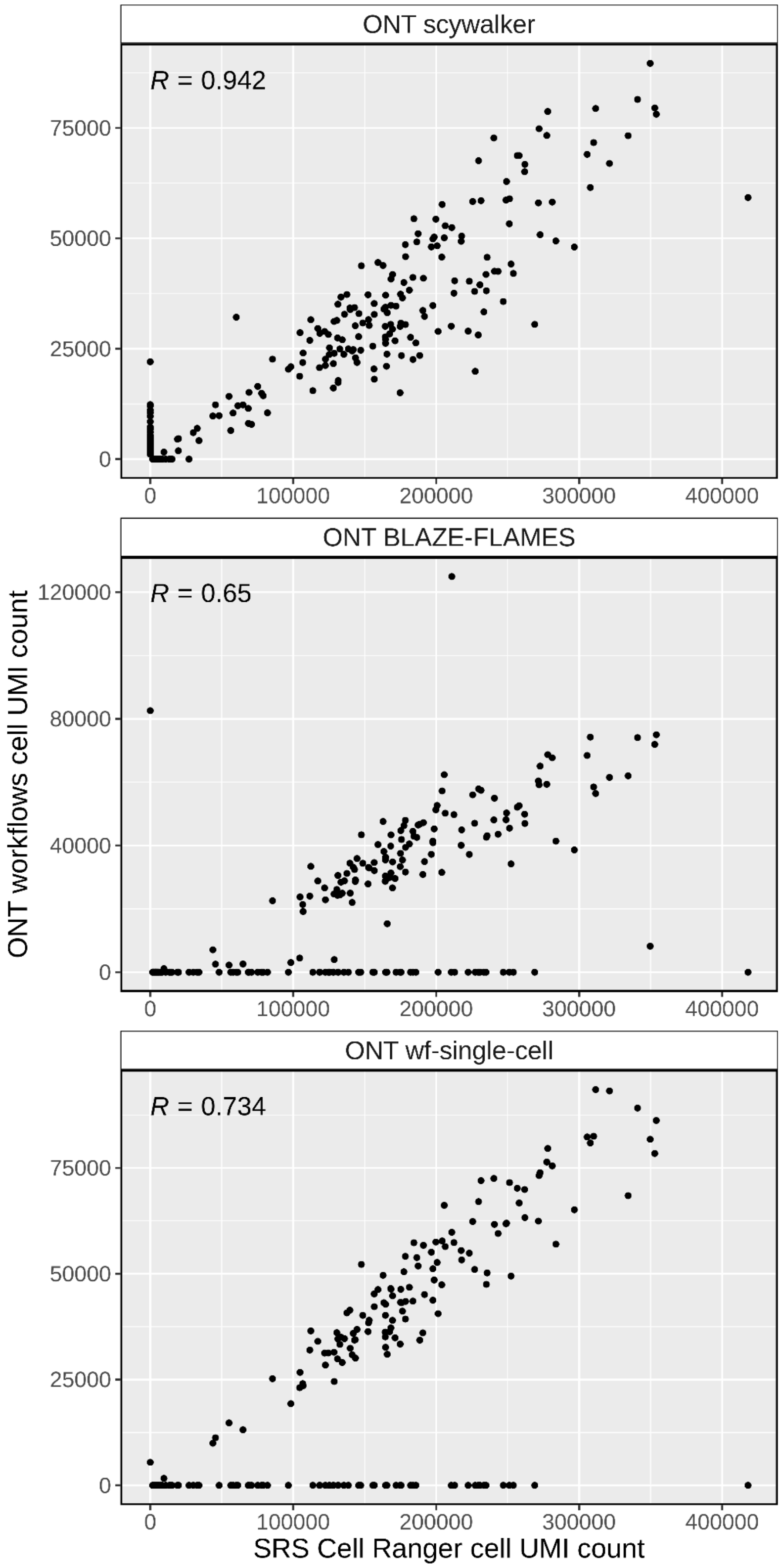


Supplementary Fig. 2: ONT scywalker, BLAZE-FLAMES, and wf-single-cell pipelines derived UMI counts per cell compared to their respective short read Cell Ranger results for the scmixology2 data set. Workflow-specific Pearson correlation coefficients (R) are shown on the upper left corners of each panel. *y* axis, ONT pipeline UMI counts per cell based on long-read sequencing data; *x* axis, Cell Ranger UMI counts per cell based on short-read sequencing data. SRS, short-read sequencing; UMI, unique molecular identifier.

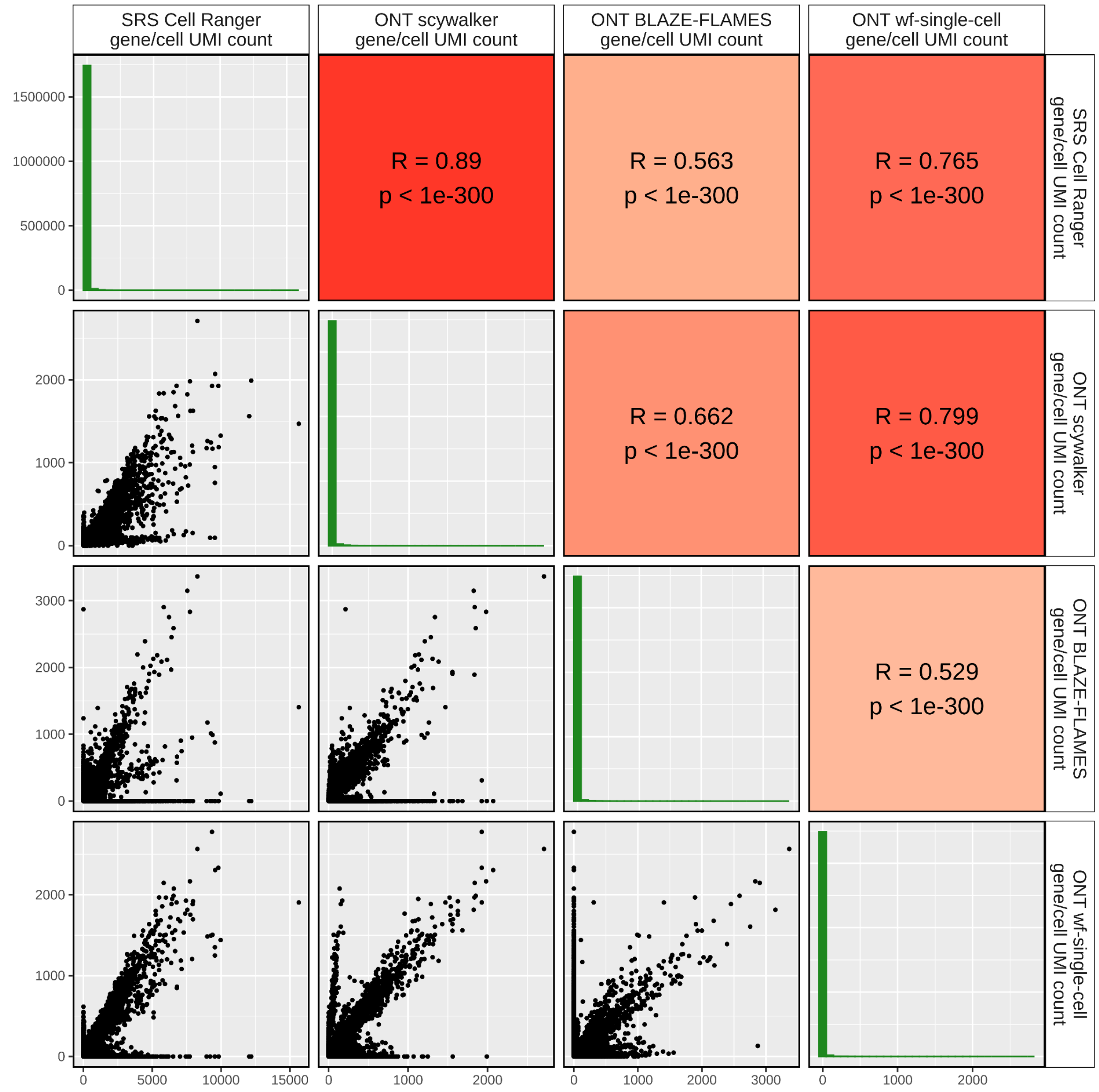


Supplementary Fig. 3: The pairs comparison plot of the UMI counts per gene and cell on the scmixology2 data set using different short and long-read analysis pipelines. The lower part of the matrix shows the scatterplots of the UMI counts per gene and cell derived from different pipelines, the diagonal part of the matrix shows the green histogram bins (n=30) of these counts, while the upper part of the matrix shows the Pearson correlation coefficients (R) and p-values (p) of correlations. A color gradient is used for the upper part of the matrix based on correlation coefficients, where higher coefficients are displayed in a darker red color. SRS, short-read sequencing; UMI, unique molecular identifier.

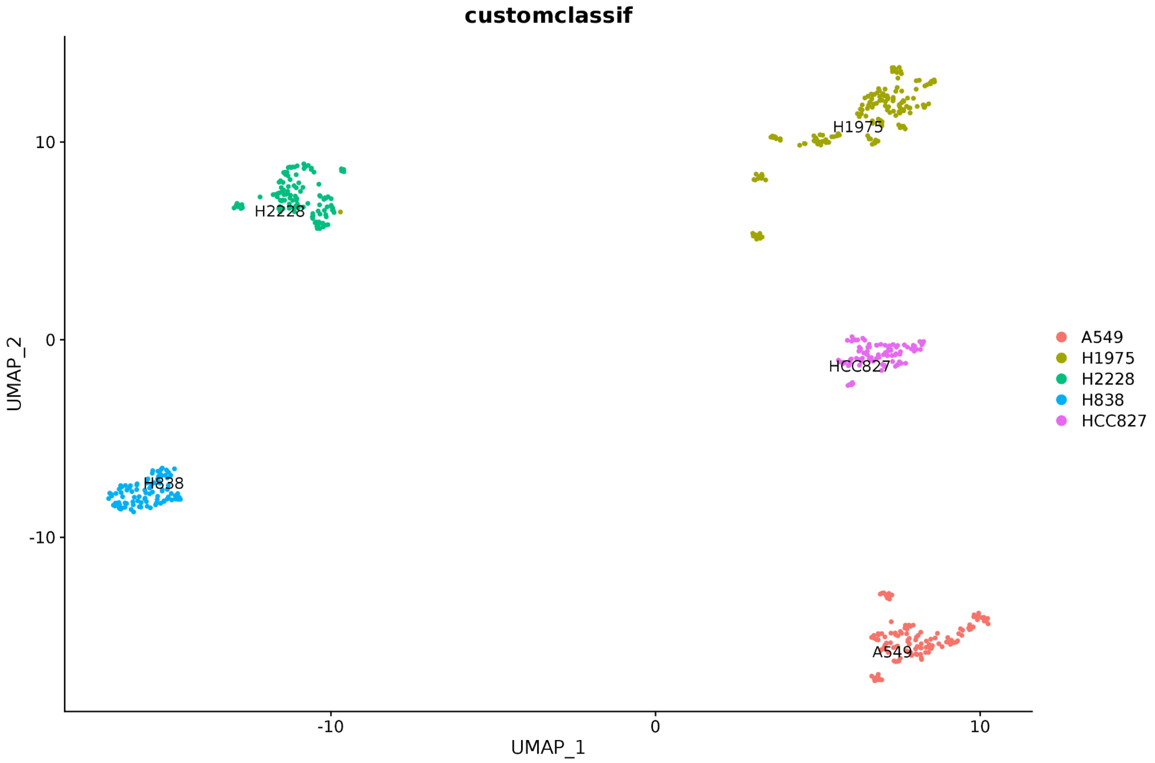


Supplementary Fig. 4: UMAP plot of scywalker results for the scmixology2 data set. Cell types predicted by ScType using a custom marker set for this experiment are indicated in color

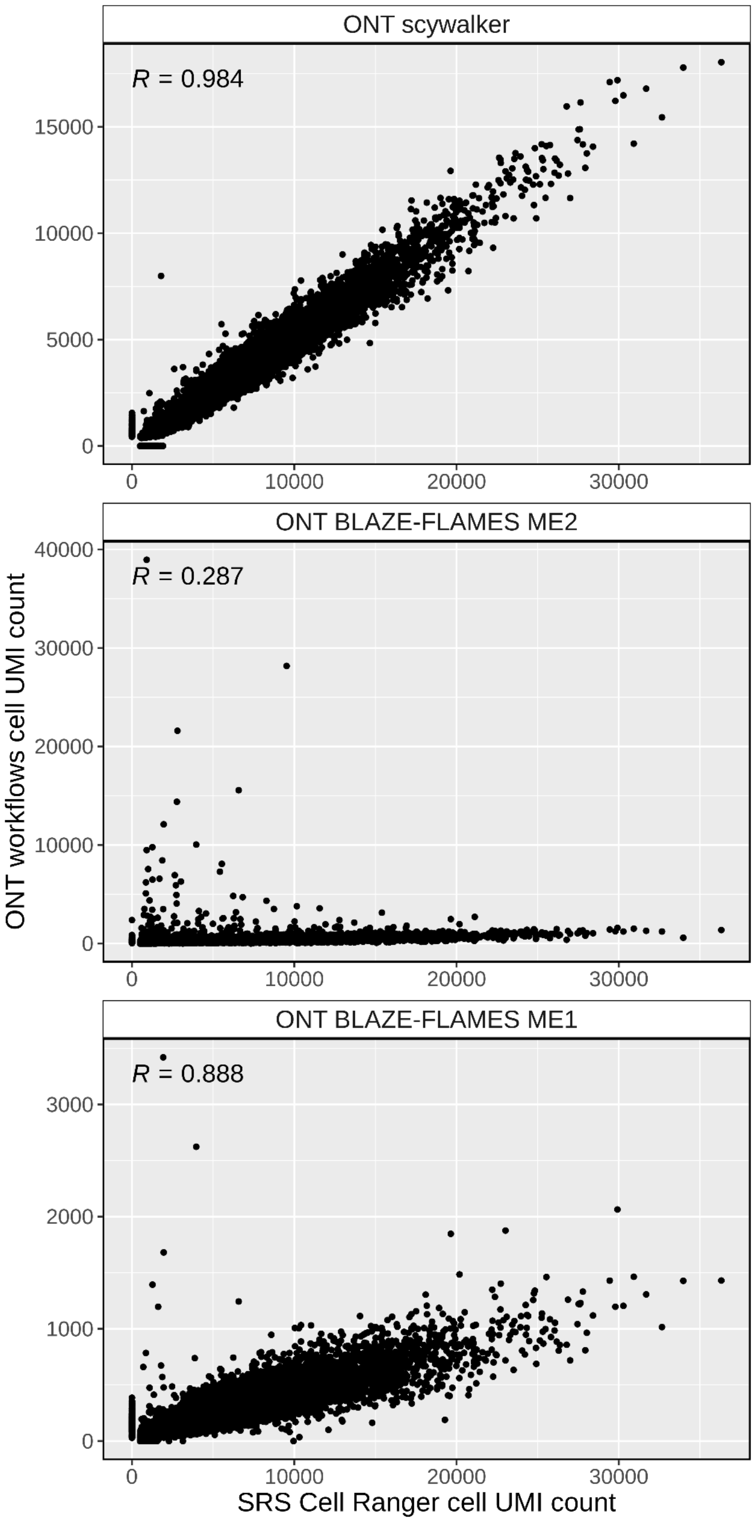


Supplementary Fig. 5: ONT scywalker and BLAZE-FLAMES pipelines derived UMI counts per cell compared to their respective short read Cell Ranger results for the brain1 data set. BLAZE-FLAMES pipeline was run with both max edit distance settings of 2 (ME2) and 1 (ME1). Workflow-specific Pearson correlation coefficients (R) are shown on the upper left corners of each panel. *y* axis, ONT pipeline UMI counts per cell based on long-read sequencing data; *x* axis, Cell Ranger UMI counts per cell based on short-read sequencing data. SRS, short-read sequencing; UMI, unique molecular identifier; ME, max edit.

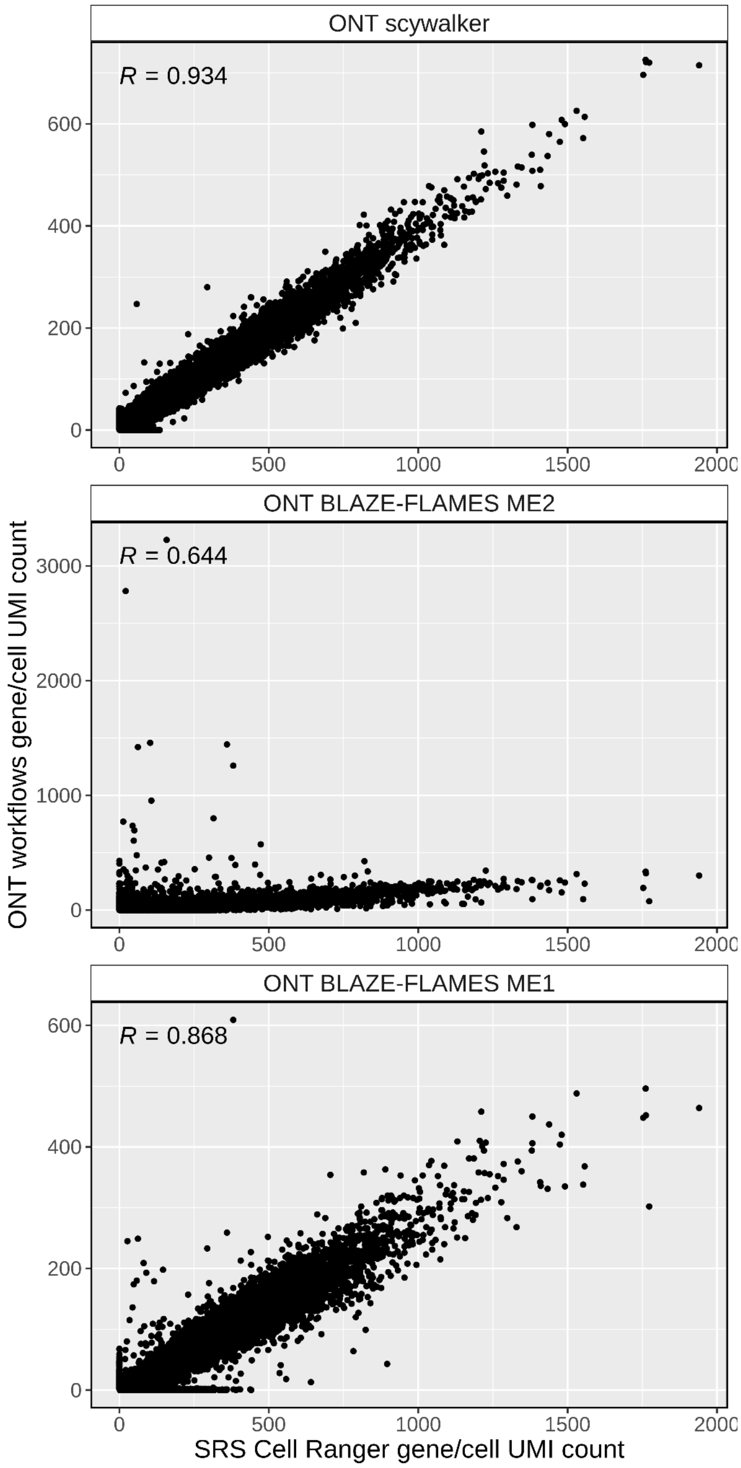


Supplementary Fig. 6: ONT scywalker and BLAZE-FLAMES pipelines derived UMI counts per gene and cell compared to their respective short read Cell Ranger results for the brain1 data set. BLAZE-FLAMES pipeline was run with both max edit distance settings of 2 (ME2) and 1 (ME1). Workflow-specific Pearson correlation coefficients (R) are shown on the upper left corners of each panel. *y* axis, ONT pipeline UMI counts per gene and cell based on long-read sequencing data; *x* axis, Cell Ranger UMI counts per gene and cell based on short-read sequencing data. SRS, short-read sequencing; UMI, unique molecular identifier; ME, max edit.

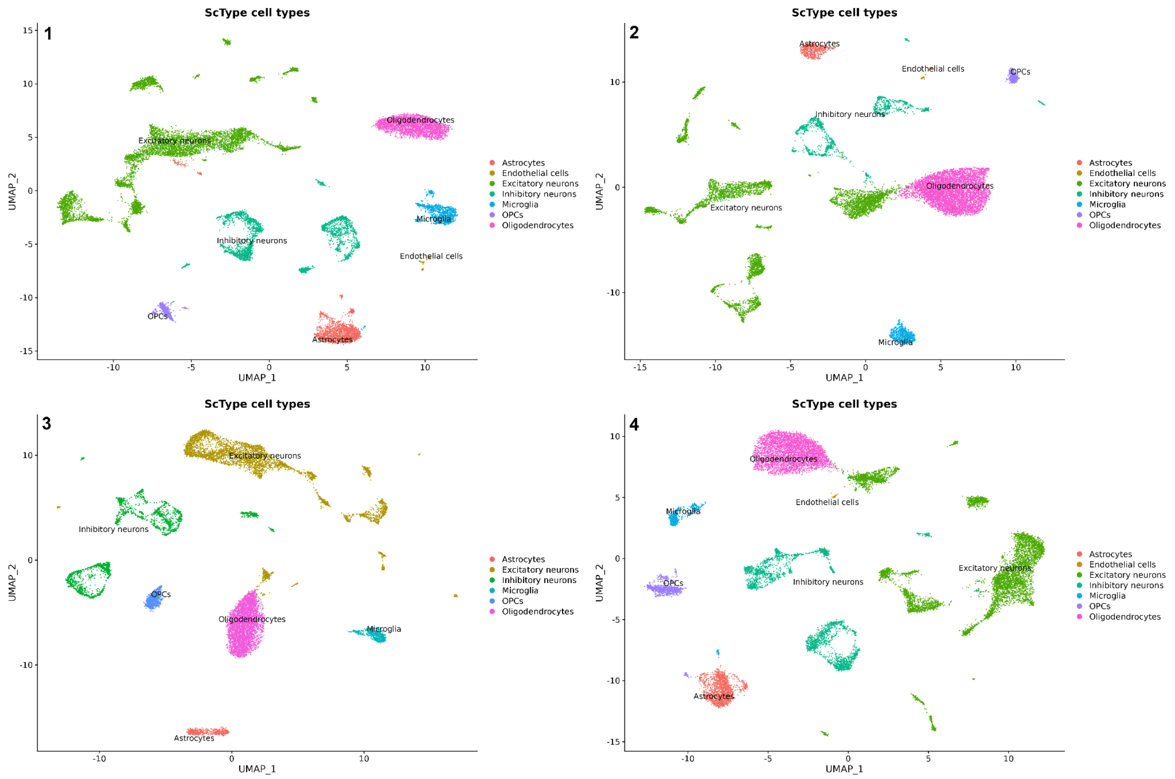


Supplementary Fig. 7: UMAP plots generated by scywalker showing cell-type assignments by ScType in different colors for brain 1-4.

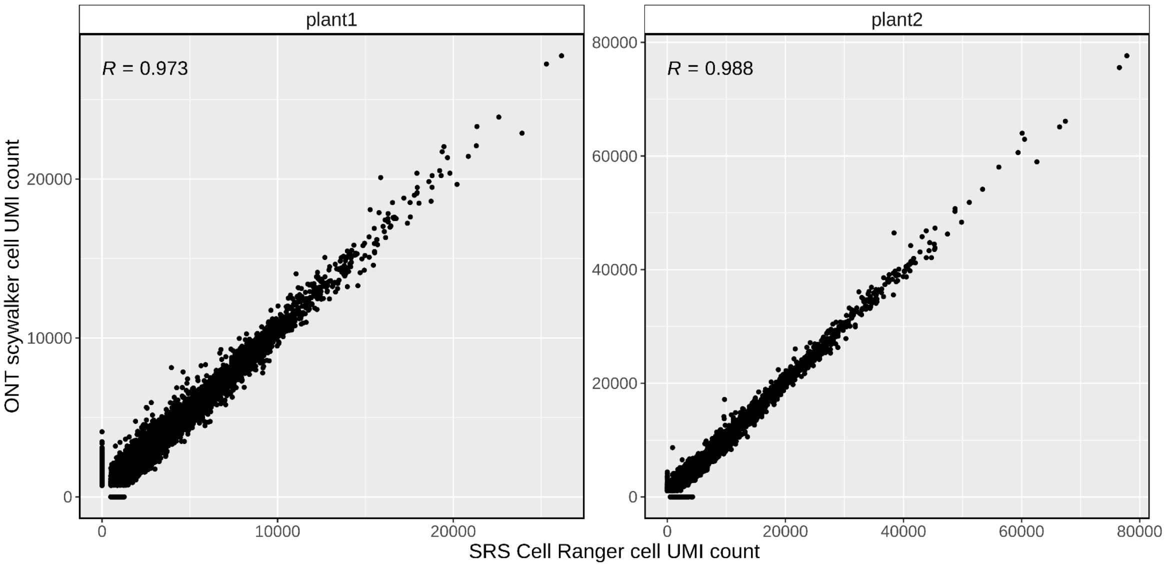


Supplementary Fig. 8: Scywalker UMI counts per cell compared to their respective short read Cell Ranger results for the two plant samples.Sample-specific Pearson correlation coefficients (R) are shown on the upper left corners of each panel. *y-axis*, scywalker UMI counts per cell based on long-read sequencing data; *x-axis*, Cell Ranger UMI counts per cell based on short-read sequencing data. SRS, short-read sequencing; UMI, unique molecular identifier.

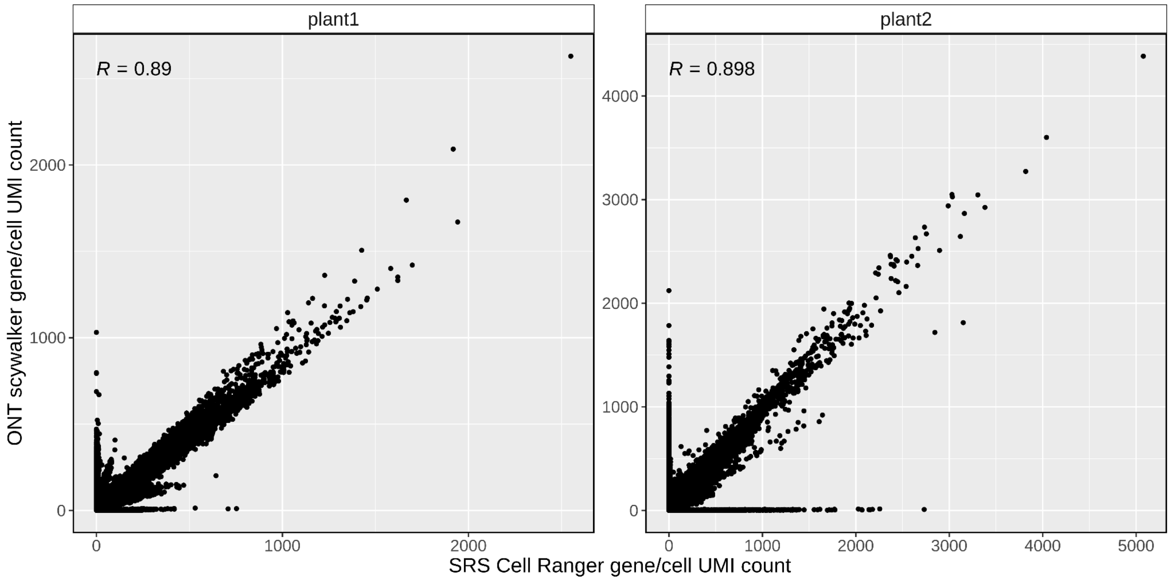


Supplementary Fig. 9: Scywalker UMI counts per gene and cell compared to their respective short-read Cell Ranger results for the two plant samples. Sample-specific Pearson correlation coefficients (R) are shown on the upper left corners of each panel. *y-axis*, scywalker UMI counts per gene and cell-based on long-read sequencing data; *x-axis*, Cell Ranger UMI counts per gene and cell-based on short-read sequencing data. SRS, short-read sequencing; UMI, unique molecular identifier.

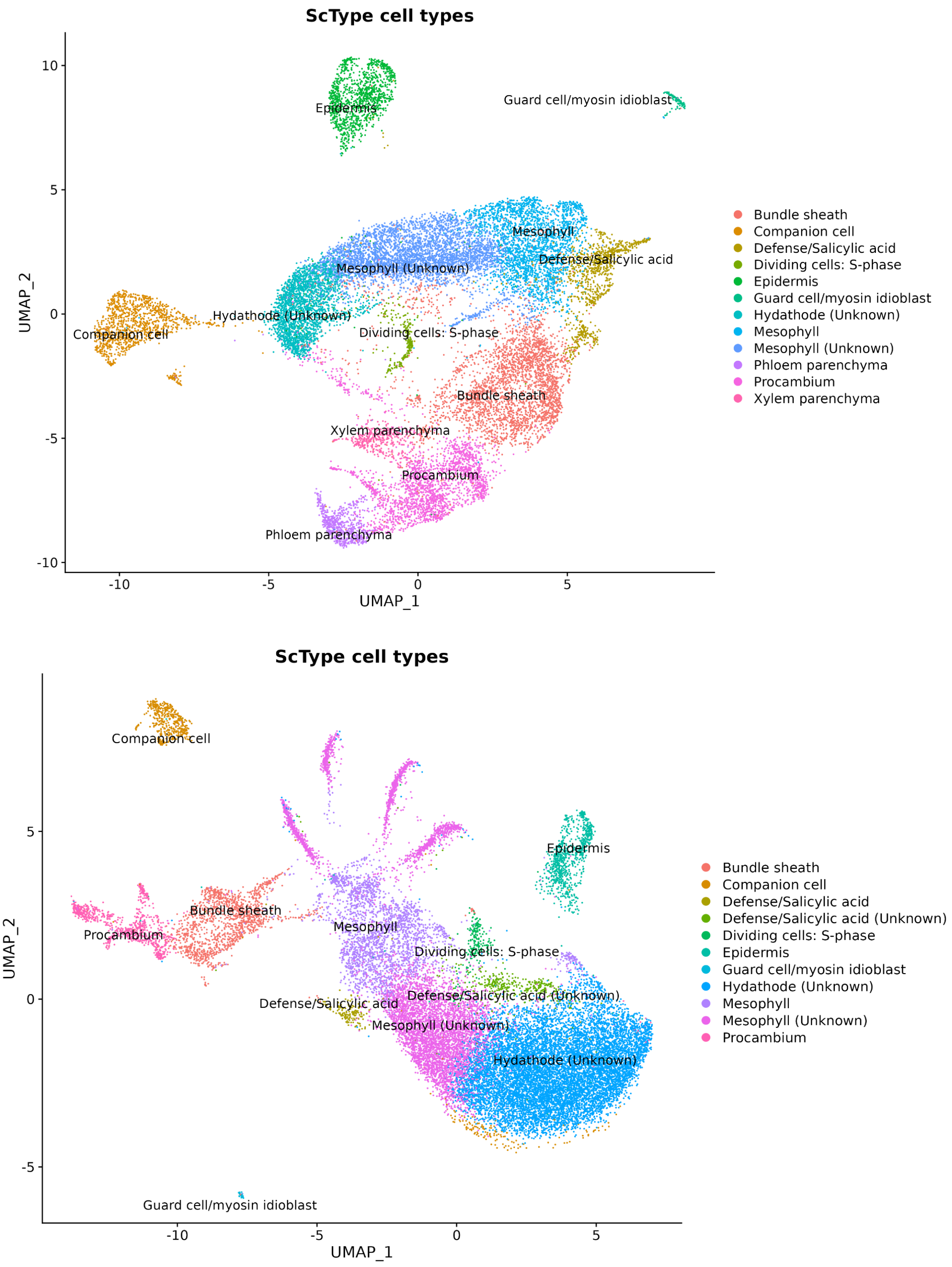


Supplementary Fig. 10: UMAP plot generated by scywalker showing cell-type assignments by ScType in different colors for plant sample 1 and 2.

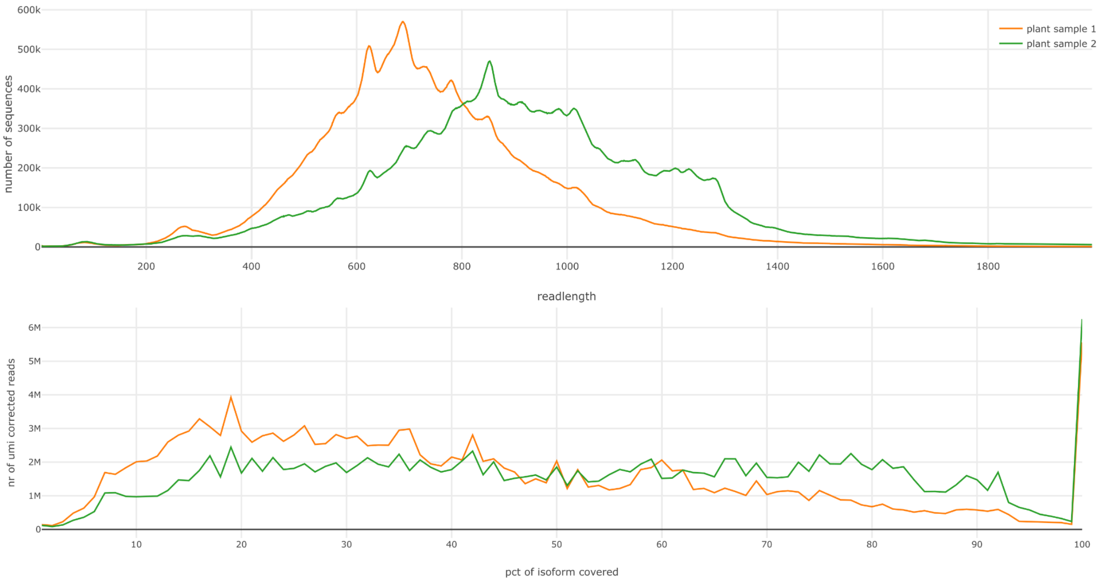


Supplementary Fig. 11 Effect of read length on coverage. The top panel shows the distribution of read lengths for plant samples 1 and 2. The bottom panel shows the number of reads covering a certain percentage of their isoform.
